# From Transporter to Motor: Evolutionary and Structural Insights into the Emergence of Prestin’s Area-Motor Activity in Mammals

**DOI:** 10.1101/2025.04.11.648445

**Authors:** Nicolás Fuentes-Ugarte, Tiaren Ruiz-Rojas, Felipe Garcia-Olave, Alvaro Ruiz-Fernandez, Jose Antonio Garate, Victor Castro-Fernandez, Raul Araya-Secchi

## Abstract

Prestin, a member of the SLC26A family, is essential for the electromotility of mammalian outer hair cells, converting voltage changes into mechanical work. In contrast, nonmammalian orthologues function as anion transporters. To investigate the molecular and structural basis of this functional divergence, we performed ancestral sequence reconstruction (ASR) of prestin across vertebrates, followed by structural modeling using AlphaFold2-multimer and molecular dynamics simulations. We identified more than 200 amino acid substitutions along the lineage that lead to placental mammals, with early substitutions concentrated in the transmembrane domain (TMD) and late substitutions clustering in the STAS domain, particularly in the intervening sequence (IVS). Structural modeling and simulation revealed that early substitutions modulate protein–lipid interactions and interhelical contacts. In placental mammals, the IVS-loop adopts a distinct conformation that places a negatively charged patch near the chloride access pathway, potentially affecting the ion dynamics and voltage responsiveness. These structural transitions occurred without major rearrangements of the global fold of prestin, supporting a notion in which the novel function evolved through distributed substitutions within a conserved scaffold. Our findings illustrate how molecular exaptation, and incremental structural remodeling enabled the repurposing of an ancestral anion transporter into a voltage-sensitive area-motor, providing a framework for understanding the molecular evolution of complex biophysical traits central to auditory neuroscience.

## INTRODUCTION

The auditory system of mammals exhibits remarkable sensitivity and frequency selectivity. These features rely on the cochlear amplifier, a mechanism that enhances sound-induced vibrations within the cochlea to improve sensitivity and frequency resolution. At the heart of this amplification process are outer hair cells (OHCs), specialized sensory cells that exhibit electromotility (EM), the ability to rapidly change body length in response to electrical stimulation[1, 2, 3, 4, 5]. EM is driven by the area-motor protein prestin, densely packed in the lateral membrane of OHCs[3, 6, 7].

Prestin is a member of the SLC26A family, which predominantly comprises anion transporters[8]. As such, nonmammalian prestin orthologs function as anion exchangers[9, 10, 11], while mammalian prestin has evolved into a direct voltage-to-displacement converter, operating on millisecond or faster timescales, which are essential for high-frequency sound amplification[3, 5, 7, 12, 13].

Phylogenetic analyzes have revealed that mammalian prestin evolved from an anion transporter present in a pre-mammalian ancestor[14, 15, 16]. Mammalian prestin exhibits minimal transport capacity and pronounced nonlinear capacitance (NLC), a hallmark of voltage sensitivity[11, 17, 18, 19]. Furthermore, an inverse relationship has been established between anion transport and motor function, suggesting that transport capacity was lost as voltage-dependent NLC was gained [10, 15]. Thus, the unique function of mammalian prestin is the result of evolutionary adaptations that have repurposed an ancestral anion transporter into a highly specialized motor protein. This functional shift from ion transport to electromotility remains incompletely understood, particularly with regard to how key substitutions have fine-tuned this mechanism in placental mammals.

Considerable efforts have been devoted to identify the regions and residues where the differences between mammalian prestin and nonmammalian prestin are most pronounced and critical for the generation of NLC and EM. Early work identified specific segments within the transmembrane domain (TMD), notably TM1–2 and TM9–11, as essential to NLC/EM and determined that replacing these regions in the zefrafish prestin by those of gerbil, a synthetic prestin capable of transport and NLC could be constructed[20]. Others showed that the extracellular loop between TM3 and TM4 (EC-loop) is also essential for NLC/EM[21, 22, 23]. Furthermore, increased flexibility of TM6 was proposed to contribute to NLC/EM[24], as well as certain conserved and non-conserved cysteines[25]. In addition, membrane thickness has been shown to affect mammalian and nonmammalian prestin differently[26].

Recently, the structures of human(HsPres)[27], dolphin(TtPres)[24], and gerbil (MuPres)[28] prestin have provided invaluable information on its functional organization and its interactions with the lipid membrane. From these structures, molecular dynamics (MD) simulation studies have shown the existence of elevator-like movements as part of its conformational change cycle[29] and how prestin structure affects the organization of the membrane in its surroundings[30]. However, fundamental questions remain regarding how voltage triggers conformational change[24, 31, 32], and how disordered regions, absent in experimental structures such as the IVS-loop, contribute to the specialized function of mammalian prestin[33].

Here, we address these gaps employing ancestral sequence reconstruction (ASR) combined with structural modeling, conformational sampling and atomistic molecular dynamics (MD) simulations to identify and contextualize key substitutions in prestin evolution. We map these changes to prestin domains and regions and then use conformational sampling with AlphaFold2-multimer (AF2-multimer)[34, 35] and MD simulations to evaluate their structural effects. This approach not only corroborates the relevance of known functional sites, but unveils previously unexplored features, notably in the IVS-loop of the STAS domain. By modeling nonmammalian prestin structures for the first time, we offer an evolutionary perspective on how incremental substitutions collectively reshaped an ancestral anionic transporter into the high-speed area-motor protein essential for mammalian hearing.

Our results indicate that early substitutions alter contacts between residues on the TMD and between the protein and the surrounding lipids, which point toward a gradual reduction in transport capacity along with an enhanced voltage sensitivity. Despite these functional changes, the TMD architecture remains largely unchanged until more punctuated substitutions appear after the divergence between mammals and reptiles. These later substitutions cluster in the EC-loop and, more strikingly, in the IVS of the STAS domain, giving rise to pronounced structural differences between nonmammalian and mammalian prestin. In particular, alterations in the IVS-loop result in the emergence, in placental mammals, of a patch of negatively charged residues (negative patch or NP) that could influence the access/exit of chloride ions from the binding site, suggesting a pivotal role for this region in the unique electromechanical behavior of mammalian prestin.

## RESULTS AND DISCUSSION

### Ancestral Sequence Reconstruction Identifies Critical Evolutionary Substitutions in Prestin Domains

We constructed a phylogenetic tree for the chordate SLC26A5 subfamily as shown in Figure 1 A (for a full version of the tree, see Supplementary File 1), identifying five monophyletic groups (fish, amphibians, reptiles, birds and mammals) with a consistent topology compared to previous studies [36, 37]. Using ancestral sequence reconstruction (ASR)[38], we inferred 12 ancestral sequences, listed in Table I, with high confidence (mean posterior probability > 0.95; see also Supplementary Figures S1, S2, S3 and S4).

**Figure 1.**
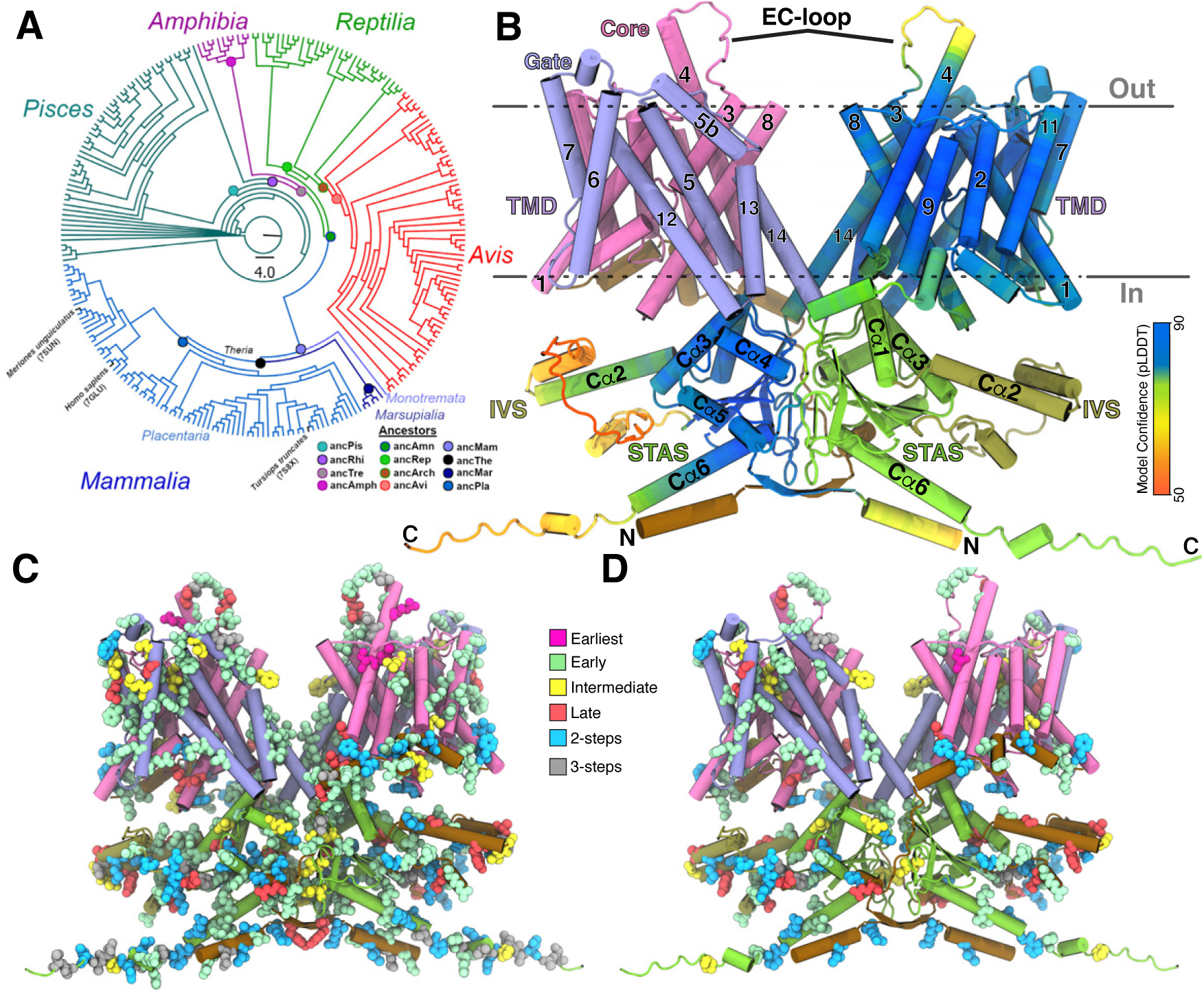
Ancestral sequence reconstruction of SLC26A5 members. (A) Maximum likelihood phylogenetic tree of prestin (SLC26A5) from Chordata. Sequences colored as follow: Dark cyan Pisces, dark magenta Amphibian, green Reptilian, red Birds, light blue monotremata mammalians, navy blue marsupialia mammalians and bright navy blue placentaria mammalians. The extant sequences of *Homo sapiens (HsPres)*, *Meriones unguiculatus (MuPres)* and *Tursiops truncates (TtPres)* are highlighted with their respective PDB accession codes. Circles indicate nodes where ancestral sequences were inferred in this study, while the scale bar represent the mean number of substitutions per site. (B) Structural model of full-lenght HsPres. Protein shown in cartoon representation with the monomer on the right labeled by AF2-multimerV3 confidence score (pLDDT) and the other by domain (N-terminal domain, ochre; core domain, pink; gate domain, violet: STAS domain, light-green and IVS, tan. Dashed black lines depict the approximate position of the membrane (C-D) Mapping of the substitutions onto the HsPres model showing (C) all and (D) non-conservative substitutions. Domains colored as in B and substitutions colored as: earliest, magenta; early, lime; intermediate, light-yellow; late, red; 2-step light-blue; and 3-step, gray.

**Table I.** Robustness of the Ancestral Sequence Reconstructions.

| Ancestor | Description | Node Support | ASR Mean Posterior Probability ± SD |
| --- | --- | --- | --- |
| ancPis | Last common ancestor from Pisces | 99.98% | 0.974 ± 0.092 |
| ancRhi | Last common ancestor from Rhipidistia | 96.68% | 0.972 ± 0.095 |
| ancTre | Last common ancestor from Tetrapoda | 99.98% | 0.952 ± 0.125 |
| ancAmp | Last common ancestor from Amphibia | 77.94% | 0.973 ± 0.090 |
| ancAmn | Last common ancestor from Amniota | 99.98% | 0.974 ± 0.088 |
| ancRep | Last common ancestor from Reptilia | 93.41% | 0.977 ± 0.083 |
| ancArch | Last common ancestor from Archosauria | 99.98% | 0.978 ± 0.081 |
| ancAvi | Last common ancestor from Avis | 99.98% | 0.976 ± 0.084 |
| ancMam | Last common ancestor from Mammalia | 99.98% | 0.959 ± 0.108 |
| ancThe | Last common ancestor from Theria | 99.98% | 0.962 ± 0.113 |
| ancMar | Last common ancestor from Marsupialia | 99.98% | 0.996 ± 0.033 |
| ancPla | Last common ancestor from Placentaria | 99.98% | 0.999 ± 0.001 |

Our analysis focused on the evolutionary pathway from fish to placental mammalians (ancPis → ancRhi → ancTre → ancAmn → ancMam → ancThe → ancPla; following the nomenclature introduced in Table I) using as initial reference (outgroup) the closest homologous transporter, human SLC26A6 (HsA6), and human SLC26A5 (HsPres) as a model of current placental mammalian prestin. For the evolutionary analysis, we also included the AltALL sequences for ancPis, ancRhi, ancTre, ancAmn, ancMam, and ancThe, as described in the Materials and Methods section. The substitution and sequence identity analyzes along the evolutionary trajectory were performed using the most probable ancestral sequences; however, in the substitution analysis, uncertain positions where an alternative state with a posterior probability >0.2 may exist are indicated (see Tables S3, S4, S5, S6 and in the Supplementary File 2). Sequence identity was high within groups (90% in nonmammals, 80% in mammals), and dropped to 60% between them (Supplementary Table S1).

We identified 206 substitutions along this evolutionary pathway: 42 in the N-terminal domain, 82 in the transmembrane domain (TMD) and 82 in the sulfate transporter and antisigma (STAS) domain. Substitutions were classified as early (ancAmn → ancMam; n=107), intermediate (ancMam → ancThe; n=19), or late (ancThe → ancPla; n=21), plus four earlier substitutions (ancTre → ancAmn). We also found substitutions showing two (n = 38) and three steps (n=17) in the transition from ancAmn to ancPla (or HsPres) that are indicative of regions with greater variability (the complete list of substitutions is presented in the Supplementary Tables S3, S4, S5, S6 and in the Supplementary File 2). In terms of proportions, *i.e.* the percentage of substituted residues, the N-terminal domain presents a 56% of substitutions, the TMD 19% and the STAS domain 35%, suggesting that the TMD is more conserved and less prone to substitutions.

Regarding the nature of the substitutions, we found that 72% (n = 148) of the substitutions are non-conservative, i.e., they change the physical-chemical properties of the residue. To identify these substitutions, we classified the amino acids into the following groups: hydrophobic aliphatic (I, V, L), hydrophobic aromatic (F, Y, W), other hydrophobic (A, G, M), polar negative (E, D), polar positive (K, R), polar neutral (Q, N, C, S, T), and other (H) and (P)[39]. Non-conservative substitutions were defined as those involving a change between these groups, these substitutions are indicated in **bold\*** throughout the text. They correspond to 62% of the substitutions in the N-terminal domain, 76% in the TMD, and 73% in the STAS domain. According to our findings, these substitutions tend to correspond to late, 2-step, or 3-step substitutions. Interestingly, the most prevalent types of non-conservative substitutions were those that result in the addition of glycines or prolines and those that result in the replacement of an apolar for an aromatic sidechain.

In summary, our reconstruction reveals that prestin evolution involved a substantial number of amino acid substitutions, with a strong bias toward late, non-conservative changes in the STAS domain and to a lesser extent in the TMD. The prevalence of substitutions that add glycine or proline, and apolar-to-aromatic substitutions highlights potentially function-altering events that occurred during the transition from ancestral transporters to the specialized motor protein observed in placental mammals.

### Conformational Sampling Reveals that the IVS-loop of the STAS domain differentiates mammalian Prestin

Placing the substitutions in the context of prestin structure is crucial to understanding their functional relevance. Thus, we generated a full-length model of human prestin (HsPres) using AlphaFold2-multimer (AF2-multimer (V3))[34, 35] to account for the regions missing in the available mammalian prestin structures, particularly the IVS-loop. In addition, we generated structural predictions for ancestral and extant prestin representatives (Figure 2 and Supplementary Figure S5). The models closely resembled the domain-swapped dimeric structure of members of the SLC26A family (Figure 1 B) and showed high confidence, indicated by the high values of the predicted local distance difference test (pLDDT) observed for most of the well-folded part of the TMD and STAS domain (Figure 2 and Supplementary Figure S5). The flexible and disordered parts (N- and C-terminal, EC-loop, IVS-loop) show the lowest pLDTT values, indicating low confidence.

**Figure 2.**
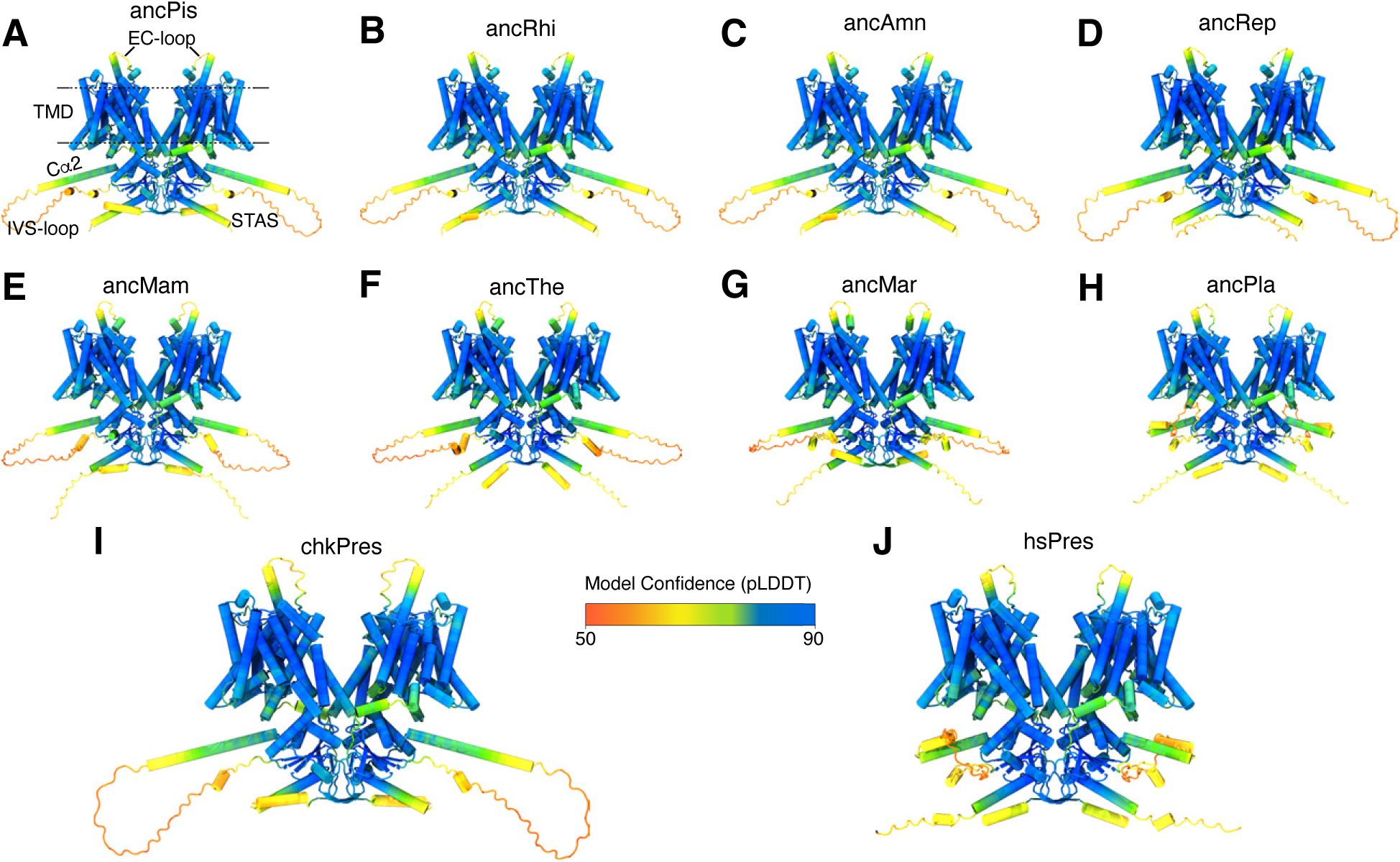
Reconstruction of ancestral prestin structures using AF2-multimer(V3). Structural predictions of ancestors from Pisces (A), Rhipidistia (B), Amniota (C), Reptilia (D), Mammalia (E), Theria (F), Marsupialia (G), Placentaria (H), chicken prestin (I) and human prestin (J) shown in cartoon representation colored by confidence (pLDDT).

Because structural predictions of flexible regions such as the EC- and IVS-loops often show low confidence, we used ensemble-based sampling to explore their conformational variability. Since AF2 predictions are based on the statistical coupling of amino acid positions across evolutionary time, this framework is well suited to examine how lineage-specific substitutions could alter structural features. We used two ensemble-based approaches: (i) stochastic sampling using multiple seeds and the *dropout* option, and (ii) varying the depth of the multiple sequence alignment (MSA) to modulate the strength of coevolutionary signals (see Methods). Similar approaches have been used to explore the conformational diversity in transporters and receptors[40, 41, 42]. For the first approach, we generated 1280 models per sequence, covering HsPres, reconstructed prestin ancestors (ancAmn, ancRep, ancThe, ancMam, ancPla) and extant representatives, while for the second approach we generated 480 models per sequence for HsPres, ancPla, ancThe and ancMam.

Both strategies primarily sampled local variations in the flexible loops, particularly the EC- and IVS-loop, while the TMD and folded parts of the STAS domain remained largely unchanged. No large-scale conformational transitions corresponding to the expanded contracted (or outward-inward-open) states associated to the functional conformational cycle of these proteins were captured.. This limitation probably arises from the presence of at least one SLC26A structure in the AF2 training set, which restricts sampling to the conformational basin represented in the data [40]. It is important to mention that the resulting ensembles should be interpreted as qualitative representations of possible conformations, rather than a quantitative representation of population distributions.

To evaluate whether the differences observed in the AF2 models reflect sequence-dependent effects rather than the results of stochastic sampling, we performed equilibrium molecular dynamics simulations on selected ancestral models embedded in POPC bilayers (ancAmn, ancRep, ancMam, and HsPres). These simulations revealed consistent interhelical interactions and loop conformations, supporting that the observed changes are stable features compatible with the structures predicted with AF2-multimer (more details in the next sections).

Across all reconstructed prestin models, the transmembrane domain (TMD) exhibited high structural similarity (RMSD-Cα ≤ 3.0 Å), with variations in the TM3-TM4 loop (extracelular or EC-loop) and the TM6-TM7 loops. Similarly, for the structured part of the STAS domain. Interestingly, the only major structural divergence observed between nonmammalian and mammalian models (ancestral and extant) was in the conformation and orientation of the intervening sequence loop (IVS-loop).

In nonmammalians ancestors and extant representatives, the IVS consistently formed an extended Cα2 helix followed by a long disordered loop (Figure 2A-D and Supplementary Figure S5A-I). In contrast, mammalian ancestors and extant representatives exhibited a distinct rearrangement of this region. Early mammalians and marsupials (ancMam, ancThe, platypus, opposum, wombat) showed a shorter Cα2 helix followed by a disordered IVS-loop (Figure 2 E-F and Supplementary Figure S5 J-L) while placental mammals (ancPla, human, gerbil, dolphin, and bat) showed a broken helix (Cα2 + Cα2’) repositioning the IVS loop toward the intracellular entrance of the adjacent monomer (Figure 2 GH and Supplementary Figure S5 MH).

This consistent difference in conformation and orientation of the IVS-loop is correlated with the differences observed between sequences of nonmammalian and mammalian ancestors and between early mammalians, marsupials and placentals (Supplementary Figure S4 and Supplementary File 2). Furthermore, they are in line with the progression of prestin function from ion transport in nonmammals followed by intermediate area-motor adaptations in monotremes to robust, finely tuned electromotility in therian mammals[10, 16, 43]. Thus, the IVS-loop represents the most prominent structural difference distinguishing mammalian prestin from their ancestral and nonmammalian counterparts. We discuss the relationship between these differences and the substitutions that we found in that region in a later section of this work.

### Mapping of substitutions into the structural models of HsPres and the reconstructed ancestors

Mapping of substitutions on the HsPres model (Figures 1 C-D and 3), we observed that the early substitutions are distributed throughout the TMD and STAS domains with non-conservative early substitutions accumulating in the TM5b, TM6, EC-loop and STAS domains (Figure 1 D, Figure3, for TM5b, TM6 and EC-loop reconstruction robustness see Table S2 and Supplementary File 2). Intermediate substitutions are found mainly in TM5b, TM6 and in the STAS domain. Late substitutions are found in the EC-loop, the TM5b-TM6 loop, and in the STAS domain. Finally, two- and three-step substitutions are found in the EC-loop and STAS domains, indicating positions with the least restriction to substitutions. In the following sections, we detail our analysis for each domain separately.

### Substitutions to the TMD show clustering in the Gate Domain

Almost 20% of the residues in the TMD (n=82) are substituted during the ancAmn ancPla transition, with 55 of them corresponding to non-conservative substitutions.. In our analysis, we found that the early and intermediate substitutions are located close and far from the chloride-binding site, whereas late substitutions are positioned further away from it. (Figure 1C-D and Figure 3). This suggests that early and intermediate changes, which are widespread across the transmembrane domain (TMD) and do not necessarily affect chloride binding, may have contributed to the loss of ion transport and the emergence of voltage sensitivity. However, late substitutions, mapped to a few clustered locations, may be responsible for the fine-tuning of prestin to provide the high sensitivity and selectivity observed in placental mammals[16].

**Figure 3.**
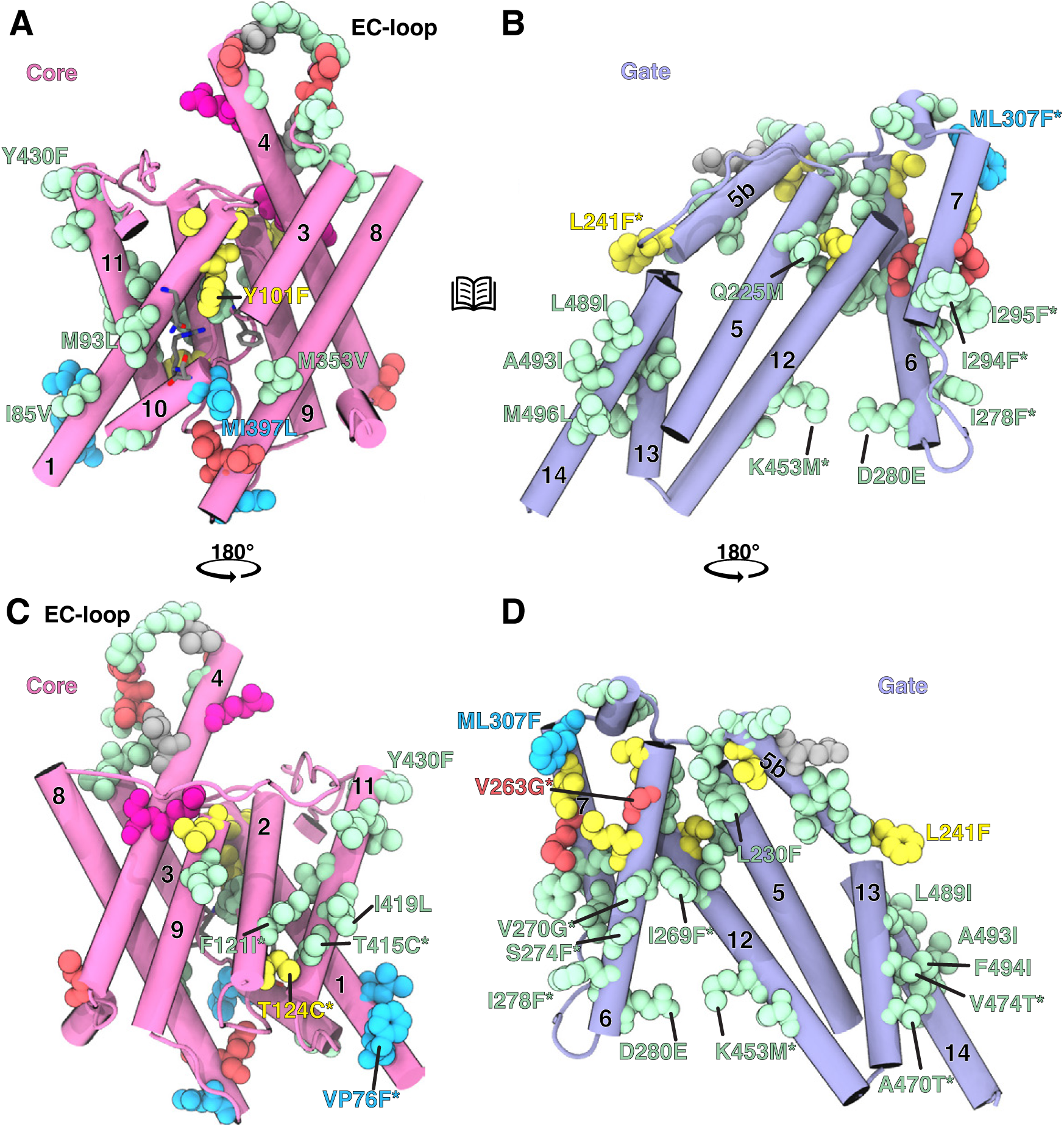
Substitutions on the TMD. (A-B) Open-book view of the core (A) and gate (B) domain interface of HsPres. (C-D) 180° rotation along the z-axis of the core domain (C) and gate domain (D) with respect to A and B respectively. In all panels the protein is shown in cartoon representation and substituted residues in VdW representation. The same color scheme as in Figure 1B-D was used. Two-step substitutions are represented as, for example, **VP76F\***, with residue numbering based on human prestin (HsPres).

Notably, not all TM helices are equally substituted, and we found more substitutions in the gate domain (scaffold) than in the core domain (transport) with the highest number of substitutions found in TM5b, TM6 and TM12 (Figure3, Table II and Supplementary Table S4). Furthermore, in both domains the majority of substitutions are located in the external face of the helix, with the exception of TM6 and TM7. In these helices, their lipid-facing region, which also forms the interface between the two, concentrates all the substitutions (Figure 3B-D).

**Table II.** Summary of TM Helices Substitutions.

| TM Helix | Subst./length | NC Substitutions | Evolutionary Step |
| --- | --- | --- | --- |
| TM1 | 8/29 | 2 | 2-step, early and intermediate |
| TM2 | 2/17 | 2 | 1 early, 1 intermediate |
| TM3 | 3/14 | 3 | All early |
| TM4 | 5/27 | 4 | 3 earliest, 1 late, 1 3-step |
| TM5 | 2/23 | 2 | Early |
| TM5b | 7/11 | 4 | 4 Early, 2 intermediate, 1 3-step |
| TM6 | 11/23 | 10 | 7 Early, 2 intermediate, 2 late |
| TM7 | 9/15 | 7 | 6 early, 1 intermediate, 1 late, 1 2-step |
| TM8 | 4/35 | 4 | 1 early, 2 late, 1 2-step |
| TM9 | 2/17 | 1 | 1 early, 1 intermediate |
| TM10 | 2/9 | 2 | 1 early, 1 2-step |
| TM11 | 7/21 | 2 | all early |
| TM12 | 5/27 | 5 | 4 early, 1 intermediate |
| TM13 | 4/15 | 4 | all early |
| TM14 | 4/20 | 3 | all early |

It has been shown that the function of prestin and other members of the SLC26A family involves changes in the orientation between the core and gate domains [24, 27], therefore, we expected to find an accumulation of substitutions in their interface. However, as can be seen in Figure 3A-B, very few substitutions are found at the core-gate interface. From the core domain, we found **M93L\*** (TM1), Y101F (TM1), **M353V\*** (TM8) and **MI397L\*** (TM10) and from the Gate domain we found only **Q225M\*** (TM5). Some of these substitutions are located near the chloride binding site, which highlights their relevance for NLC/EM. For example, in TM1 substitutions **M93L\*** and Y101F (early) and **T136P\*** (intermediate) when mutated individually from their mammalian to nonmammalian version (that is, **L93M\***, F101Y, **P136T\***) resulted in changes in the parameters of NLC and when mutated together completely abolished NLC/EM [20]. Furthermore, it has been shown that residues L139 and V353 participate in conformational rearrangements associated with motion of the core (transport) domain, consistent with elevator-like transitions that regulate exposure of the chloride-binding cavity [29].

Although the core–gate interface shows few substitutions, we identified a network of residue changes near the interdomain linkers connecting TM7–TM8 and TM11–TM12 (for TM7, TM8, TM11 and TM12 reconstruction robustness see Table S2 and Supplementary File 2). These include a combination of early, intermediate, and late substitutions located on the extracellular side of TM5, TM5b, TM6, TM7, and TM12 in the gate domain and the extracellular side of TM11 in the core domain (Figure 3) and may have changed the coupling between domains and therefore the functional conformational change.

One of the helices in the TMD with the largest number of substitutions is TM6, the majority being non-conservative. This helix has been suggested to play a crucial role in NLC/EM due to the accumulation of glycines in a unique pattern in mammalian prestin[24]. Our analysis showed that of the glycines found in placental prestin, two correspond to early substitutions and one to a late substitution (Figure 4). Furthermore, we found several substitutions in TM6 that could also be relevant for its flexibility since they participate in interactions with TM7 and TM12 (Figure 3B-D and Figure 4) and could affect the movement of TM6. Furthermore, comparison with the nonmammalian ancestor ancAmn shows that early substitutions D280E in TM6 and **K453M\*** in TM12 could perturb an electrostatic contact network between these helices (Figure 4), which would reduce the strength of the interaction between them and allow TM6 to move more freely. Other charged residues that are part of this network, in mammalian prestin, have been shown to affect NLC/EM when mutated to glutamine[44], highlighting the role of the interaction of TM6 - TM12 in prestin function.

**Figure 4.**
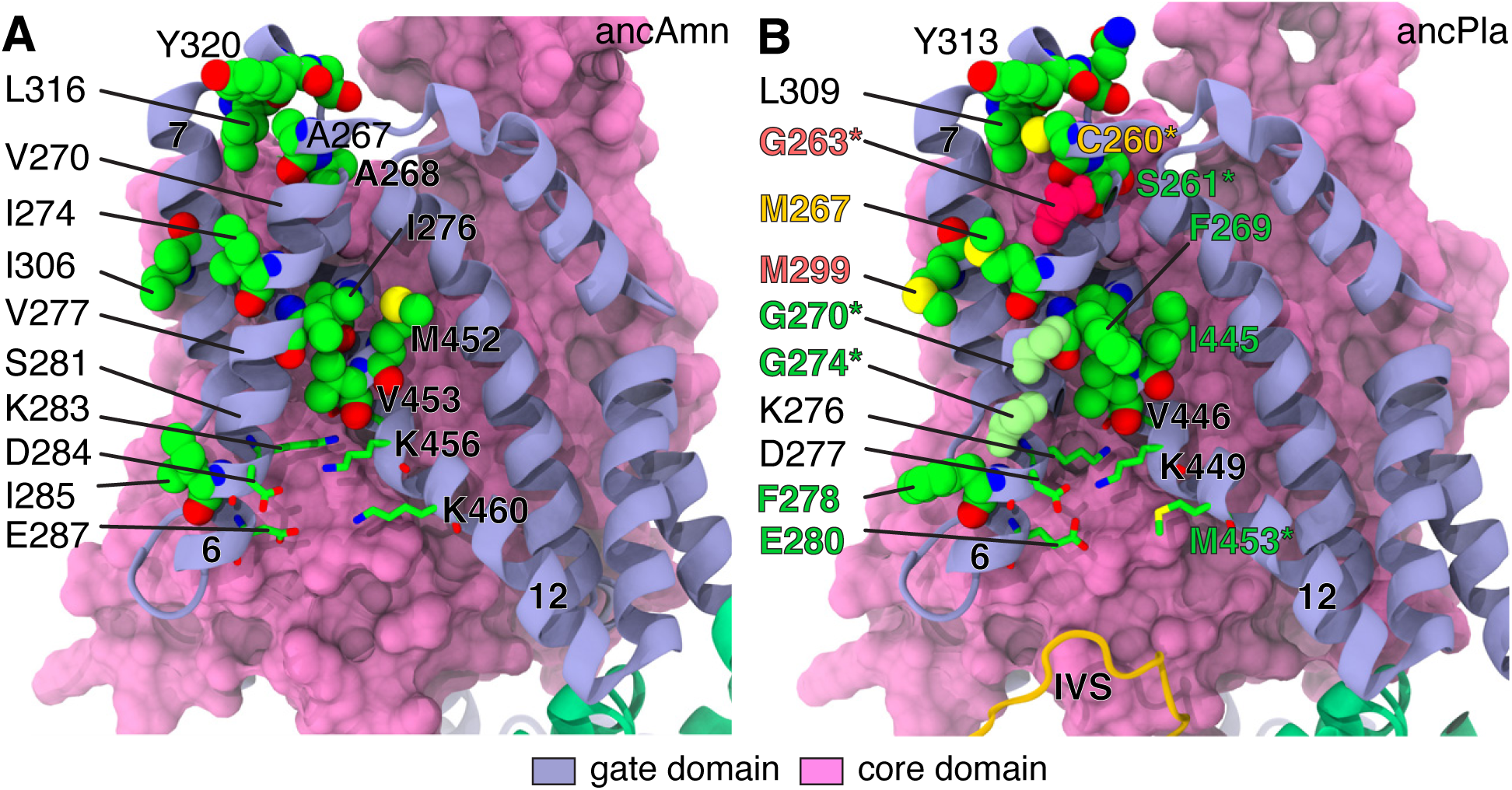
Substitutions in the TMD. (A-B) Substitutions in TM6 in the ancAmn (A) and ancPla (B) models. The gate domain is shown as ribbons and the core domain as surface colored as in Figure 1B. Substitution labels are colored as in Figure 1C and shown in VdW or licorice representation colored by element (carbon, green; oxygen, red; nitrogen, blue and sulphur, yellow). Only the TMD of one monomer is shown for clarity. Non-conservative substitutions marked with an *.

Our MD simulations revealed a slight increase in RMSF-Cα values for HsPres and ancMam compared with ancRep and ancAmn at TM6 (Supplementary Figure S6). This increase was accompanied by a broader distance distribution between TM6 and TM12, particularly between residues D277 or E280 (TM6, early) and K449 (TM12) (Supplementary Figure S7), suggesting that TM6 has greater flexibility and freedom to move in mammals. Residues D277 and E280 (and their equivalents in ancAmn and ancRep) form salt bridges primarily with K276 and R281 (both from TM6), and to a lesser extent with K449, in both mammalian and nonmammalian systems. However, the frequency of salt bridge formation between D277-K449 (8.2%) and E280-K449 (9.3%) was higher than in nonmammalians (ancAmn and ancRep), where equivalent interactions occurred at frequencies below 3%. This pattern is consistent with the bimodal distance distributions observed for these residue pairs in the nonmammalian simulations (Supplementary Figure S7A–B) and with the broader COM-COM distance distribution between TM6 and TM12 (Supplementary Figure S7C).

Together, these observations build upon the notion that the increased flexibility of TM6 in mammalian prestin results from the accumulation of glycines[24]. Our results suggest that additional substitutions in TM6, TM7, and TM12 may also contribute to enhanced flexibility and freedom of movement of TM6 by altering interhelical interactions. Consistent with our results, a recent cryo-EM study shows that TM6 and TM7 bend significantly upon membrane thinning, acting as a mechanosensitive ‘elbow’[45]. The presence of substitutions in these helices in the early, intermediate, and late evolutionary stages implies further structural modulation after the emergence of motor function, possibly reflecting the fine-tuning to the frequency range characteristic of placental mammals[16].

### Several Substitutions in the TMD affect the protein-lipid interfase

Mammalian prestin is known to be sensitive to changes in membrane tension [46, 47, 48], thickness [26, 49] and cholesterol concentration[50, 51, 52]. Notably, most substitutions identified in the TMD are located on the lipid-facing side of the helices, particularly in the gate domain (Figure 3). This suggests that the evolutionary transition from transporter to area-motor may have involved significant changes in how prestin interacts with the membrane. Interestingly, previous studies have identified differences in the response to changes in membrane thickness between mammalian and nonmammalian prestin [26, 49], and recently, the first direct comparison of structures of lizard and dolphin prestin solved with lipids of different lengths revealed that dolphin prestin undergoes a stable conformational change with membrane thinning, while lizard prestin remains in an unchanged state[**?**]. These results lend support to our observations that the conversion from a transporter to and area-motor involved changes in the protein-membrane interactions.

We identified two sets of non-conservative substitutions that may be the most relevant: The first set comprises seven substitutions in which an apolar residue is replaced by phenylalanine. They are mainly located near the extracellular or intracellular end of an helix and concentrate in the gate domain (TM5b, TM6, and TM7) with only two found in the core domain (TM1 and TM11) (Figure3). The second set of substitutions comprised four early non-conservative substitutions in TM13 that result in a unique pattern found only in mammalian prestin (**V470T\***, **V474T\***, **A475T\*** and **A478S\***). In our mammalian models, as in the available mammalian prestin structures, residues V474, S478, and T479 (residue numbering according to HsPres) are located on the exterior face of the helix, while T475 faces inward towards TM5. This results in three polar residues placed inside the lipid bilayer in the intermonomer region (Figure5A-B). These residues could use their side chain hydroxyl groups to form intrahelix (TM13) and interhelix (TM13-TM14) hydrogen bonds, a characteristic that has previously been observed when these types of residue are located in transmembrane helices [53]. Furthermore, the presence of polar residues such as Ser or Thr in the lipid bilayer has been shown to affect the thickness of the membrane and the organization of lipids around the protein [53]. More importantly, these residues increase the polarity in the intermonomer region, facilitating the interaction with the poplar headgroups of lipids or cholesterol.

Our MD simulations of ancMam and HsPres revealed stable intrahelical hydrogen bonds involving these residues and the main-chain oxygens of neighboring residues of TM13. For example, T470–K466, T474–T470, T475–I471, S478–T474, and T479–T475 form hydrogen bonds for more than 60% of the simulation time. In addition, we observed transient hydrogen bonds between the T474 side chain and the carbonyl oxygen atoms of POPC molecules.

We also detected differences in the intermonomer contacts between TM13 and TM14 (Figure 5C–F, for TM13 and TM14 reconstruction robustness see Table S2 and Supplementary File 2). In mammalian systems (HsPres and ancMam), the contact interface between the two TM14 helices begins near the middle of the helices, deep within the lipid bilayer (Figure 5C–D), where residues I494 and L497 of both monomers are closely packed. In contrast, in ancRep and ancAmn, the interaction begins closer to the edge of the lower leaflet (Figure 5E–F), particularly in ancAmn. This shift suggests a reduction in the intermonomer space in mammalian prestin. The measurement of lipid occupancy in this region (see Methods) showed a slight decrease in the average number of lipids for ancMam (14.08 ± 1.66) and a more pronounced reduction for HsPres (12.72 ± 1.79) compared with ancRep (15.22 ± 1.40) and ancAmn (14.97 ± 1.43) which may suggest an effect of these substitutions on the volume of the inter-monomer space. However, more detailed computational and experimental studies will be required to properly address these observations.

**Figure 5.**
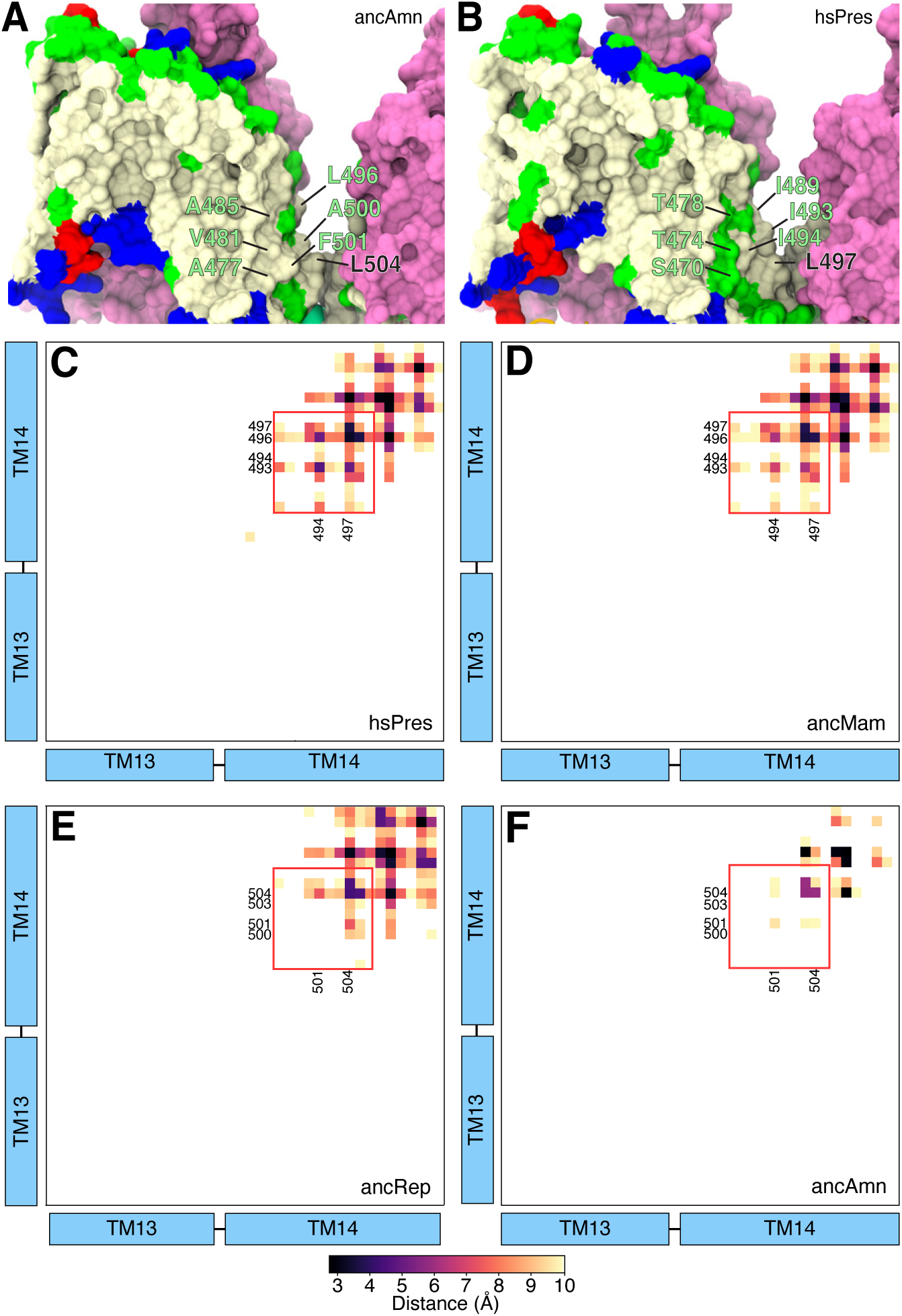
Substitutions on the TM13-14. (A-B) (A-B) Surface representation with the core domain colored as in Figure 1B and the gate domain colored by residue type (hydrophobic = white, polar = green, acidic = red, basic = blue). (A) ancAmn (B) HsPres. The same color scheme as in Figure 1 (C) was used to label the substitutions. (C-F) Average intermonomer distance between TM13-TM14 obtained from the MD-simulations of HsPres (C), ancMam (D), ancRep (E) and ancAmn (F).

In summary, we found several substitutions (mostly early) that suggest that one of the initial changes in mammalian prestin is related to its interaction with surrounding lipids. It is well known that substitutions in transmembrane helices that replace an apolar residue with a phenylalanine or a threonine can affect the disposition of the TM helix or result in remodeling of protein-lipid interactions, membrane embedding and orientation, and helix-helix interactions and dynamics[54, 55, 56]. Furthermore, these changes not only alter the protein itself but also cause the surrounding lipid and protein environment to adapt, affecting protein stability, conformational equilibrium, and interactions. Thus, these substitutions could be associated with the membrane deformation around human prestin and its lipid-mediated organization[27, 30] and the differences in response between mammalian and non-mamalian prestin to changes in membrane thickness[26].

### EC-loop configurations correlate with functional differences between nonmammalian and mammalian prestin

In the TMD we also find the extracellular loop (EC-loop), which joins TM3 and TM4 (residues 151 to 170 in HsPres). It has been shown to provide pendrin (SLC26A4) with NLC / EM when its EC-loop is replaced by that of gerbil prestin[21], while other studies have proposed that it plays a key role in EM[22, 23]. We map five early, three late, and one 3-step substitutions in this region, a remarkable accumulation for a 20 residue loop (Figure 6A). As expected, substitutions resulting in a PGG motif produce a different conformation and orientation of the loop between nonmammalian and mammalian ancestors (Figure 6). In nonmammalian ancestors, the loop faces the extracellular surface of its monomer, facilitating the formation of interactions with residues from TM4, the TM5-TM5b loop, and TM5b. In mammalian ancestors, the loop points outwards, decoupling TM3 and the loop from other regions, a situation that is more marked for the placental ancestor ancPla and HsPres.

**Figure 6.**
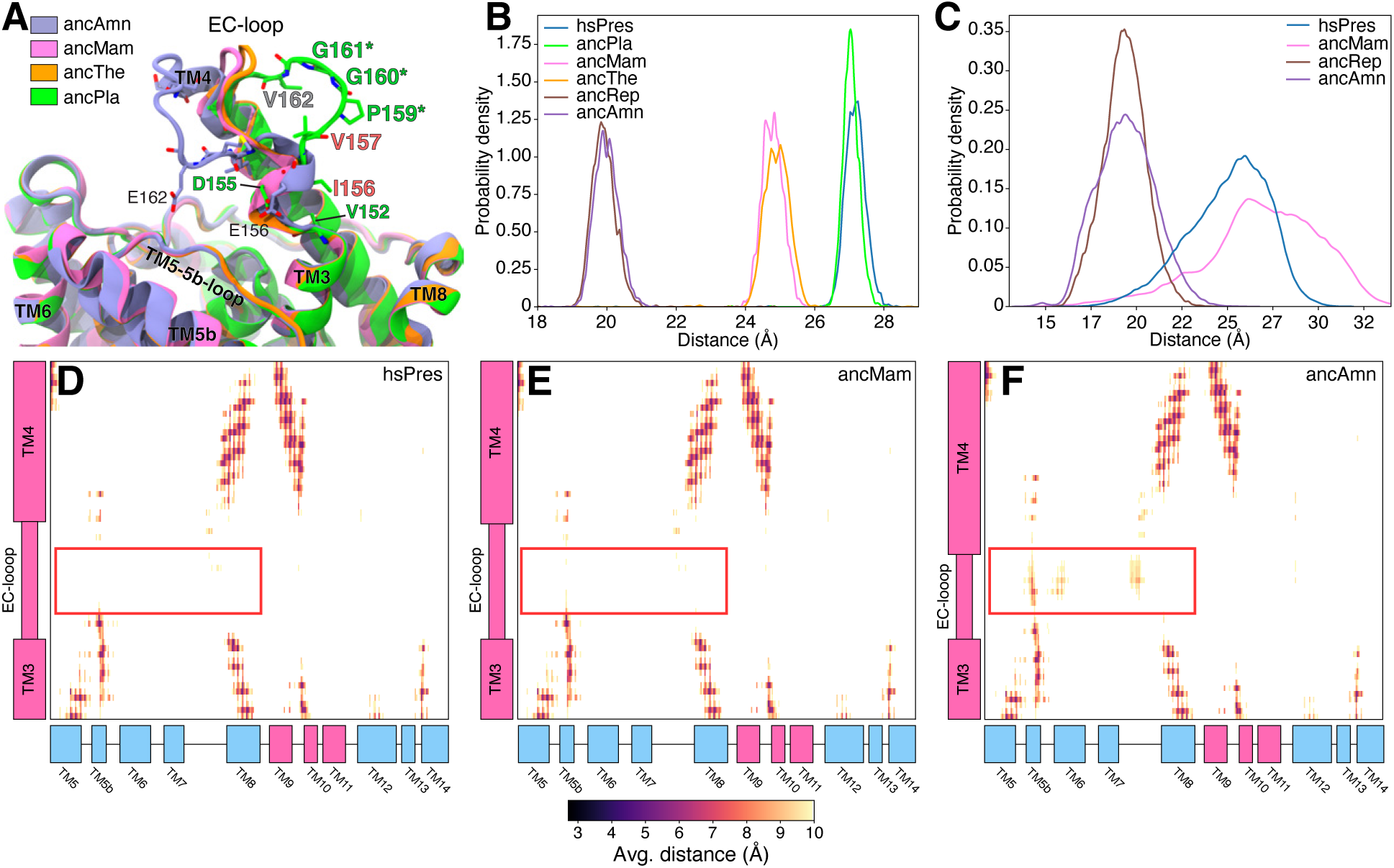
Structural effect of substitutions in the EC-loop. (A) Substitutions in the EC-loop. Proteins shown as ribbons. Substitution labels colored as in Figure 1B. Non-conservative substitutions marked with a *. (B) Distribution of the distance between the Center of Mass (COM) of the EC-loop and the COM TM5b obtained from the models AF2-conformational sampling models (C) Distribution of the distance between the Center of Mass (COM) of the EC-loop and the COM TM5b obtained from the MD simulations. (D-F) Average distance maps between TM3-TM4 and the rest of the TMD (excl. TM1-2) obtained from the MD simulations of HsPres (D), ancMam (E) and ancAmn (F).

Conformational sampling further highlights these differences, In Figure 6B-C, we show the distributions of the distance between the center of mass (COM) of the EC-loop and the COM of TM5b measured from the AF2-multimerV3 models (B) and MD simulations (C). For the AF2-multimerV3 models we observed three distinct distance distributions: nonmammalians (ancAmn, ancRep, average distance 20 Å), early-mammalians (ancMam, ancThe, average distance 25 Å) and placentals (ancPla, HsPres, 27 Å). Similarly, MD simulations show for ancAmn and ancRep 19.04±1.6 Å and 19.5±0.8 Å respectively while ancMam and HsPres simulations show average distances of 25.0±2.2 Å and 26.9±3.2 Å respectively with a broad distribution. The measurement of RMSF-Cα (Supplementary Figure S9) suggests that the wide distribution is caused by an increase in the flexibility of the mammalian EC-loop. Analysis of contacts and interactions between the EC-loop and TMD revealed differences in the contact pattern between nonmammalians and mammalians (Figure 6D-F), with the EC-loop making contact with residues of TM5b, TM6 and TM8 only in simulated nonmammalians, as shown for ancAmn in Figure 6E (red box). Replacement of a negatively charged residue (E162 in ancAmn) by a glycine (G161 in mammalians) not only affects the conformational properties of the loop, but also disrupts ionic interactions formed by it. In ancAmn and ancRep simulations, we observed that E162 consistently formed salt bridges with residues K242 and R243 (K235 and R236 in mammalian prestin, respectively) located in the loop between TM5 and TM5b.

Taken together, our ensembles of AF2 models and MD simulations suggest that substitutions in the EC-loop may influence its conformational flexibility and its interactions with neighboring regions of the transmembrane domain, particularly by reducing the coupling with the TM5–TM5b loop and TM5b. These findings remain qualitative, as they are derived from modeled and simulated systems, but point to potential structural effects that could be tested experimentally in a more detailed way than they currently have. Interestingly, TM5b and the TM5–TM5b loop exhibit several early substitutions whose structural and functional roles remain unexplored, making them promising subjects for future investigations.

### A Unique IVS-loop in Placental Mammalian Prestin results from accumulation of substitutions

The cytoplasmic STAS domain is a structural hallmark of the SLC26A family, and since the publication of the first structure[57] it has been considered an important factor for TMD movement during functional conformational change. In this domain, the greatest accumulation of substitutions is found in the intervening sequence (IVS) that connects Cα1 to sheet Cβ4 (residues 561 to 614 according to HsPres) and contains helix Cα1 (Figure 7B-C). This segment is highly variable across the SLC26A family, both in length and in amino acid composition[8]. In this section, we focus on the effect of substitutions on the structure and dynamics of this region. For details and for more details about other substitutions found on the STAS domain, see the Supplementary Discussion.

**Figure 7.**
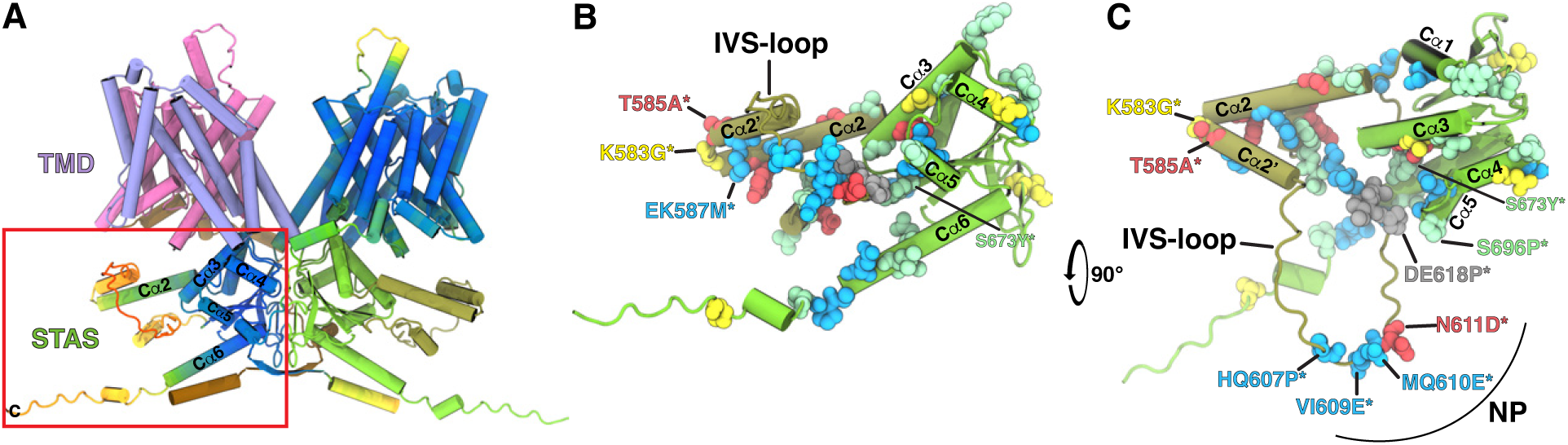
Substitutions in the IVS. (A) Schematic representation of the HsPres model highlighting with a red box the STAS domain shown in detail in panels B and C. (B-C) Detail of the STAS domain of HsPres showing only the non-conservative substitutions, highlighting substitutions found in the IVS-loop and the negative patch (NP). (B) Side view and (C) Top view (90° rotation of B along the x-axis). In all panels the protein is shown in cartoon representation and substituted residues in VdW representation. The same color scheme as in Figure 1 (BD) was used. In the IVS there is a flexible loop (IVS-loop) that connects helix Cα2 with sheet Cβ4 (residues 580 to 614 according to HsPres) that is unresolved in all currently available prestin structures and in most of the members of the SLC26A family. The only exception is a set of SLC26A3 cryo-EM structures (PDB-ID: 7XUH, 7XUJ, 7XUL, and 8IET) in which the IVS loop is resolved and placed near the intracellular entry of the adjacent monomer (Supplementary Figure S10). Unfortunately, there is no publication associated with these structures. Despite its variability, it has received limited attention in studies of the function and regulation of prestin and other members of the family. Some reports suggest potential roles in protein-protein interactions with regulators such as calmodulin[58, 59] or CTFR [60, 61]. One study found that mutations in positively and negatively charged residues within the mammalian prestin IVS loop caused modest changes in voltage operating parameters[33]. More recently, the IVS-loop in SLC26A9 has been implicated in modulating intracellular chloride dynamics[62].

In this region, we found 30 substitutions that result in clear sequence differences between nonmammalian, early-mammalian and placental ancestors (Figure 8A and Supplementary Figure S4), which include the elimination of a positively charged patch (PP in Figure 8A) and the appearance of a negatively charged patch in placental mammals (NP in Figure 8A). These differences also translate into structural differences in our models. Mammalian ancestors (ancThe onward) present the non-conservative intermediate substitution **K583G\*** which result in a broken Cα2 in two helical segments: Cα2 and Cα2’ as observed in the ancPla and HsPres models (Figure 2G-H,J, Figure 8B, Supplementary Figure S11B-E) and other mammalian representatives; (Supplementary Figure S5M-U). This characteristic is more pronounced in the ancPla and HsPres models (see Figure 8B), perhaps due to substitutions at positions 584 (3-step), 585 (late), and 587 (2-step), resulting in the NANMANA motif that has a higher helix propensity. After Cα2’, we find (P607) (two-step) and, more notably, three two-step substitutions that replace hydrophobic or polar residues with negatively charged amino acids (E609, E610, and D611) (Figure7 and Supplementary Figure S11) increasing the number of negatively charged residues resulting in a negative patch (NP) formed by residues E609, E610, and D611 (Figure 8 A). This combination of substitutions results in the placement of the NP near the entrance of the intracellular (IC) cavity of the adjacent monomer (Figure 8B-D). The NP is followed by the PIVIKSTF motif, unique to placentals, and originates from a series of two- and three-step substitutions. This motif forms contacts with residues from helix Cα3 and with a group of substitutions in helix Cα5 (P696 (early non-conservative, S to P), A697 (two-step, T to S to A), L698 (two-step, T to V to L)) that increase the hydrophobicity of this helix (Figure 7 and Supplementary Figure S11). Thus, changes that lead to the formation of Cα2’ and NP are accompanied by changes in other regions of the STAS domain that could stabilize this unique conformation.

**Figure 8.**
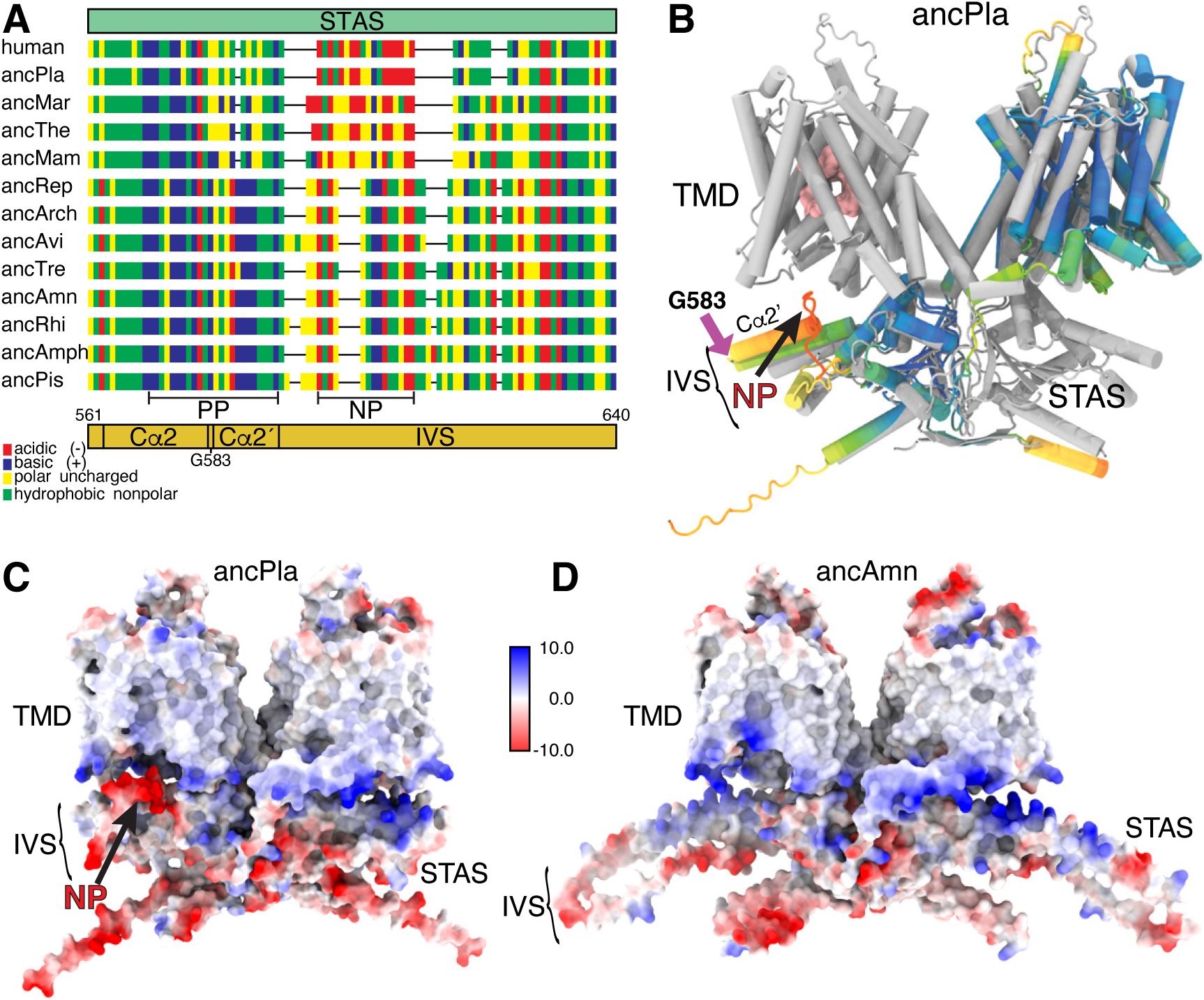
Detail of the unique features of the placental mammal IVS. (A) Fingerprint sequence alignment of the reconstructed ancestors and HsPres. Residues colored by hydropathy (acidic, red; basic, blue, polar uncharged, yellow and hydrophobic nonpolar, green). NP and PP refer to negative and positive patch, respectively. (B) Superposition of a monomer of the ancPla model (colored by pLDDT as in Figure 2) onto the structure of human prestin 7LGU[27] (grey) shown in cartoon representation. (C-D) Electrostatic potential maps of ancPla (B) and ancAmn (C).

Conformational sampling showed that nonmammalian ancestors present a continuous and extended Cα2 positioning the IVS-loop far from the intracellular (IC) entrance of the adjacent monomer. In contrast, mammalian models, especially those representing placental mammals (ancPla and HsPres), show a distinct break in Cα 2, allowing the IVS-loop to adopt conformations closer to or farther from the entrance of the IC cavity (Figure S12). To quantify this, we measured the distance between the center of mass (COM) of the IVS-loop and the COM of the chloride binding site of the adjacent monomer (Figure 9A-B) and the helical content in the IVS of the models (Figure S13A). Our results identified two distinct structural states in placental mammals, a “close” and a “far” state (bimodal distribution), compared to early-mammalian and nonmammalian models that predominantly retain a single “far” state characterized by a continuous Cα2 helix. Our findings suggest that intermediate and late substitutions significantly altered the IVS-loop in mammalian ancestors, introducing a structural hinge at residue G583 (Figure 9 B) and a unique negatively charged patch (NP) close to the chloride entry/exit point of the adjacent monomer. Although previous studies have indicated that these charged residues slightly influence nonlinear capacitance (NLC) [33], their precise functional role remains unclear. However, the distinct conformation of the placental IVS-loop raises the possibility that it could modulate chloride ion dynamics by restricting ion exit or trapping ions near the binding site, potentially influencing voltage sensing or allosteric coupling - an idea supported by recent findings in SLC26A9 that point in a similar direction[62].

**Figure 9.**
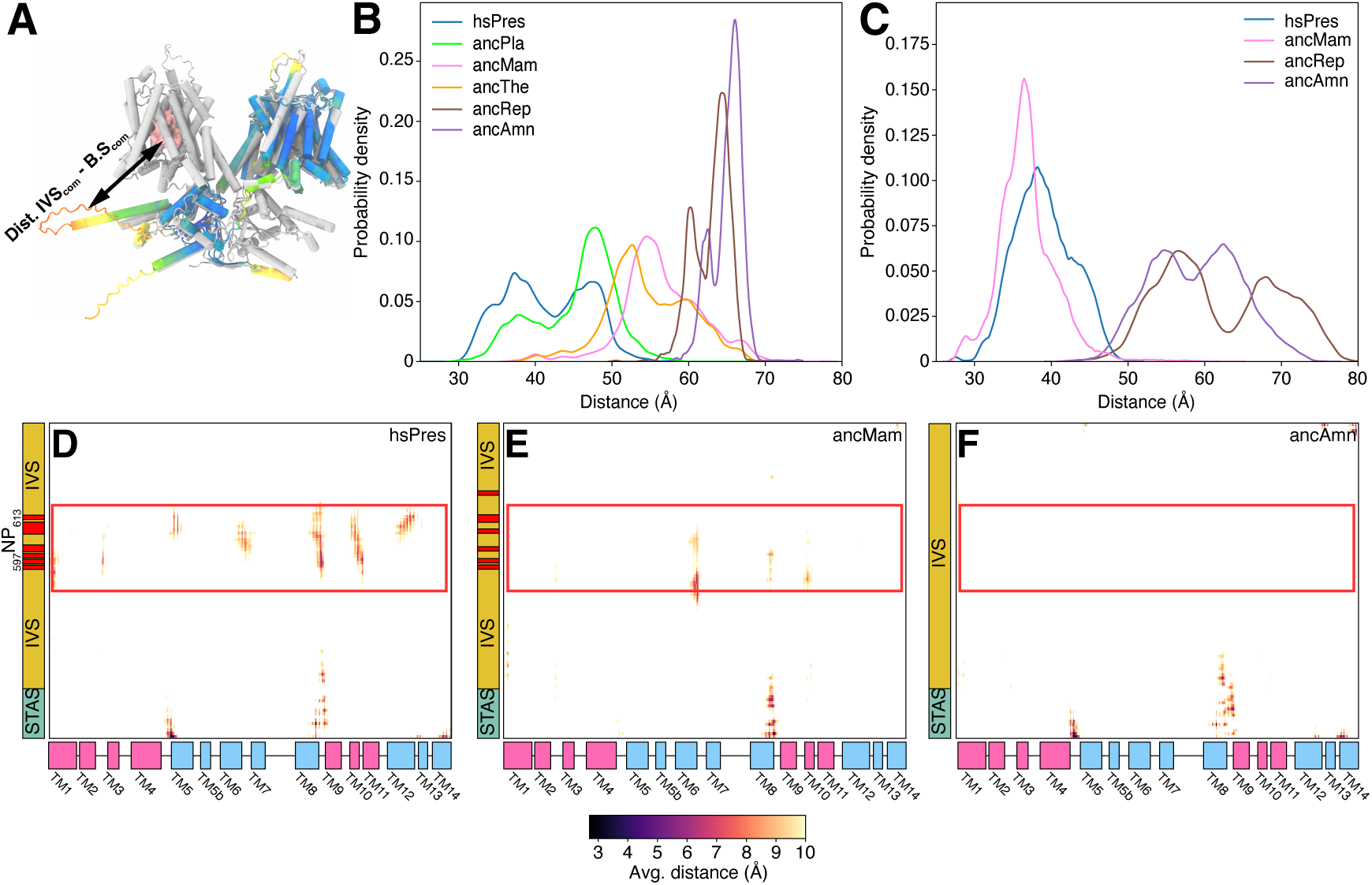
Conformational analysis of the IVS-loop. (A) Schematic representation of the distance between the COM of the IVS-loop of one monomer and the chloride-binding site of the adjacent monomer. (B) Distribution of the distance between the COM the IVS-loop of one monomer and the chloride-binding site of the adjacent monomer obtained from the models sampled with AF2-multimer(V3). (C) Distribution of the distance between the center of mass of the IVS-loop of one monomer and the chloride-binding site of the adjacent monomer obtained from MD simulations. (D-F) Average distance maps between the IVS of one monomer and the TMD of the other obtained from the MD simulations of HsPres (D), ancMam (E) and ancAmn (F). The data shown correspond to the average distance between residues from STAS-A - TMD-B and STAS-B - TMD-A. Prestin topology features are colored as in Figure 1

To further assess the robustness of our structural predictions, we performed conformational sampling of extant prestin representatives, which produced a similar pattern to that observed for ancestral reconstructions (Supplementary Figure S14). We also applied MSA subsampling, a technique used to explore alternative conformational states [40, 42, 63]. Although no major conformational transitions were detected, the ‘far’ state of the IVS-loop appeared sensitive to the strength of the coevolutionary signal, suggesting that this conformation may represent an artifact and that the ‘close’ state more accurately reflects the structure of the placental IVS-loop (Supplementary Figure S15).

We next explored these structural differences through MD simulations of representative models of HsPres, ancMam, ancRep, and ancAmn. In line with the AF2-Multimer (V3) models, we measured the intermonomer distance between the COM of the IVS-loop and the COM of the Cl^−^ binding site (Figure 9C), the helical content of the IVS-loop (Supplementary Figure S13B), and the average distance between the residues of the IVS-loop and the transmembrane domain (TMD) of the adjacent subunit (Figure 9D–F). In HsPres, the IVS-loop remained close to the neighboring Cl^−^ binding site, with an average COM–COM distance of 39.0 ± 3.9 Å, in accordance to the “close state” observed in AF2-multimer conformational sampling (Figure 9B). For ancMam, despite showing a larger average distance in the AF2-multimer confromational sampling (56.95 ± 5.3 Å) from the MD simulations, we obtained an average distance of 36.9 ± 3.8 Å, very similar to HsPres (Figure 9B). In contrast, non-mammalian ancestors exhibited greater distances, with averages of 65.0 ± 1.9 Å for ancAmn, 63.0 ± 2.3 Å for ancRep (Figure 9C). Regarding the secondary structure, HsPres showed a pronounced loss of helicity at positions 582–583, followed by the formation of helix Cα2^′^ in up to 80% of the simulation time (Supplementary Figure S13B), while ancMam shows a shorter Cα2 helix that ends at position 581. AncAmn and ancRep maintain their longer Cα2 helices during the simulations. The contact maps in Figure 9D–F show the average distance between the IVS-loop of one monomer and the TMD of the other, and it is clear that for mammals, in particular HsPres, the NP is close to the cytoplasmic side of TM5, TM6, the TM8-TM9 loop, TM10, TM11 and the TM12-TM13 loop, while no contact is observed for ancAmn. Furthermore, for HsPres we observed the formation of transient salt bridges (between 5% to 30% of simulation time) formed by E608, E610, and E611 from the IVS-loop with residues from TM5 (R211), TM6 (K276 and K283), TM8 (K364), TM12 (R463), the TM12-TM13 loop (K466) and deep into the chloride binding site (S398 (TM10), K449 (TM12)). Similarly, for ancMam we found that E609 and E610 establish transient salt-bridges with R211, K283 and K364. We noticed that the negatively charged residues in the IVS-loop could interact interchangeably with the residues on the TMD which reflects the flexibility of the loop. Interestingly, none of the residues in the TMD correspond to a substitution, they are all conserved and have been reported to affect prestin NLC [32, 44]. Our distance analysis also showed interesting differences in the contact pattern between the IVS and the rest of the STAS domain. These observations show that the PIVIKSTF motif mentioned earlier establishes close contacts with residues from helices Cα3 and Cα5 of the STAS domain (Supplementary Figure S16 A), while for ancMam, and more notoriously for ancAmn, the distance between these regions is larger (Supplementary Figure S16B-C). As mentioned earlier, helices Cα3 and Cα5 also contain substitutions that could stabilize the interactions between these regions facilitating the formation of the “close” state.

Given the position of the IVS-loop in the “close” state and recent evidence suggesting that the IVS-loop of SLC26A9 may influence the dynamics of chloride ions at the binding site [62], we analyzed chloride occupancy within the binding site of our simulated models. Consistent with Cryo-EM structures, a single Cl^−^ ion typically occupied the site at a time (Supplementary Figure S17). However, in some simulations of ancAmn and ancRep, transient double or even triple occupancy events were observed. The site remained occupied for more than 70% of the simulation time in mammalian systems, with slightly lower values for ancAmn and ancRep. Estimation of residence times revealed faster ion turnover in nonmammalian systems and slower, more variable residence times in mammalian systems, as shown in Table III. These results indicate that chloride ions tend to remain trapped within the binding site for longer periods in mammalian systems. Interestingly, recent structural data indicates that a chloride ion remains bound to prestin even during conformational transitions [45], although not tested directly in that work, these results appear to be consistent with our proposal that the mammalian IVS loop creates a permanently charged environment near the binding site that could affect chloride dynamics.

**Table III.** Cl^−^ residence time.

| Group | ChainA (ns) | ChainB (ns) | A+B (ns) |
| --- | --- | --- | --- |
| ancAmn | $0.26 \pm 0.89$ | $0.19 \pm 0.48$ | $0.22 \pm 0.68$ |
| ancRep | $0.11 \pm 0.31$ | $0.13 \pm 0.23$ | $0.12 \pm 0.27$ |
| ancMam | $1.47 \pm 24.32$ | $0.65 \pm 3.74$ | $0.89 \pm 13.63$ |
| HsPres | $0.74 \pm 6.28$ | $0.58 \pm 3.94$ | $0.65 \pm 5.07$ |

A+B correspond to the average over the two monomers. Results for HsPres and
ancMam correspond to the average from 2 independent simulations.

In summary, the IVS-loop of the STAS domain stands out as a highly diverse and lineage-specific region that underwent substantial remodeling during prestin evolution. Structural and sequence changes in placental mammals, including the formation of a hinge at G583, the emergence of a negatively charged patch, and associated substitutions in neighboring helices, appear to support a unique conformation near the intracellular entrance of the adjacent protomer. Although the precise functional consequences remain to be determined, our results suggest that these changes could contribute to fine-tuning of the voltage sensitivity and chloride accessibility, highlighting the IVS-loop as a potential regulatory element in the electromechanical behavior of prestin.

Taken together, these analyses indicate that while AlphaFold2 sampling captures local flexibility, the broader architectural differences observed across the evolutionary pathway, such as altered protein-membrane interactions, changes to helix flexibility, and reorientation of loops, reflect sequence-encoded effects. The consistency of these characteristics in both the AF2 predictions and the MD trajectories reinforces the notion that the observed differences likely represent accessible states made possible by evolutionary substitutions rather than stochastic sampling within a single conformational landscape.

It is important to emphasize that the AF2-based ensemble approaches used in this work generate plausible structural hypotheses derived from evolutionary inputs and not explicit conformational populations. However, the consistent trends reproduced by MD simulations provide support to our claims that the structural variations we describe are related to evolutionary substitutions rather than to sampling artifacts. Given that AF2 uses coevolutionary correlations, it is particularly well suited to reconstruct the structural consequences of ancestral mutations, especially when complemented with other methods such as MD simulations.

### A structural and evolutionary model for the emergence of area-motor function

Our findings recapitulate the progression of prestin function from ion transport in nonmammals through intermediate area-motor adaptations in monotremes to robust, finely tuned electromotility and corresponding frequency selectivity in therian mammals by a distributed remodeling of its architecture as it transitioned from a passive anion transporter to an active area-motor protein[10, 16, 43].

Although the overall scaffold of the TMD and STAS domains remained conserved, key substitutions introduced new physicochemical properties that collectively support the voltage-driven mechanical output.

In the TMD, substitutions altered membrane-facing surfaces and interhelical contacts. Aromatic and polar residues became more prevalent at the protein-lipid interface, potentially enhancing membrane anchoring and deformation. Substitutions in the TMD also resulted in enhanced flexibility and freedom of movement, as shown for TM6 and TM12. Remarkably, recent experimental evidence provides strong support for these predictions [45].

The most striking innovations occurred in the IVS-loop. This flexible element underwent structural rearrangements in placental mammals, adopting a conformation that places a negatively charged patch near the ion access site, a feature that may directly modulate chloride access or voltage sensitivity.

Together, these changes support an evolutionary scenario in which the area-motor function emerged through fine-tuning of existing elements within a conserved scaffold. This mosaic of adaptations may be the basis for the ability of prestin to convert voltage into mechanical work with extraordinary speed and precision.

## CONCLUSIONS

We have combined evolutionary modeling and atomistic simulations to offer a link between ancestral sequence changes and emergent structural and dynamical properties. The conformational ensembles generated by AlphaFold2 provide a qualitative view of possible structural rearrangements, and the MD simulations establish that these features are stable.

Our findings provide a structural and evolutionary framework for understanding how prestin acquired its unique area-motor function in mammals. By integrating ancestral sequence reconstruction with structural modeling, simulation and comparative analysis, we reveal that the transformation of prestin from a canonical anion transporter into a voltage-sensitive motor protein involved a distributed set of molecular innovations.

Rather than arising from a single defining mutation or domain acquisition, the functional transition inferred from our analyzes may have resulted from the gradual accumulation of substitutions that modulate membrane embedding, helix packing and interdomain coupling. These include changes in lipid-facing residues within the transmembrane region and rearrangements of flexible loop segments observed in both AF2 models and MD simulations. These observations should be viewed as qualitative structural hypotheses rather than direct mechanistic evidence, pending experimental validation.

This stepwise repurposing of conserved structural elements offers a model for how novel functional traits can evolve through subtle refinement of existing architectures, illustrating how exaptation operates at the molecular level[64]. Our work highlights the broader principles of functional innovation in membrane proteins.

## METHODS

### Multiple sequence alignment, phylogenetic inference, and ancestral sequence reconstruction

For the multiple sequence alignment (MSA) of chordate prestin, sequences were retrieved from the non-redundant protein database (nr) using BLASTp, with HsPres (Uniprot accession: P58743-1) as reference. Sequences exhibiting an identity greater than 98% were filtered through CD-HIT[65], while those with lengths greater than 870 and less than 570 residues were excluded. The resulting set of sequences (n = 248, see the sequence list in Supplementary Table S7) was aligned with a structural profile derived from the CryoEM structures of HsPres ([27]; PDB-ID: 7GLU), TtPres ([24]; PDB-ID: 7S8X) and MuPres ([28];PDB-ID: 7SUN) using STAMP[66] and ClustalW[67]. Regions corresponding to loops were manually inspected to ensure consistency of secondary structure boundaries and to minimize alignment ambiguity. The evolutive model for MSA was chosen using SMS2.0 [68] through the Akaike information criterion (AIC) and the Bayes information criterion (BIC). A maximum-likelihood phylogenetic tree was built employing PhyML[69], using a JTT matrix as a fixed model, with gamma-shaped variation across sites and a proportion of invariant sites. The robustness of the nodes was assessed using the likelihood ratio test and ancestral sequences were inferred using the empirical Bayes method with PAML 4.4 [70], employing the Lazarus tool as described by Handson-Smith[71]. Probability calculations for each amino acid were calculated at each alignment position for each target node, selecting the amino acids with the highest probabilities. The positions of the gaps were inferred by maximum likelihood using a binary evolutionary traits model with a rate of change using Mesquite [72] as previously reported[73]. Specifically, the MSA was converted into a binary matrix (1 = residue, 0 = gap), and discrete ancestral states were estimated using the phylogeny derived from protein sequences. For each ancestral node, the most probable state (residue or gap) was then assigned to each position. The resulting binary reconstruction was aligned with the amino acid reconstruction, and the positions most likely to correspond to gaps (state 0) were removed from the ancestral sequences.

Given the potential bias inherent to maximum likelihood ancestral reconstructions, we additionally generated AltALL sequences. In these alternative reconstructions, the second most probable amino acid was incorporated at positions where its posterior probability exceeded 0.2, following the approach described by Eick et al.[39]

### Generation of structural models

Full-length dimeric structural predictions of ancestral prestin sequences together with human prestin (HsPres), chicken (ChkPres) and other representatives of mammalian and nonmammalian prestin (see Supplementary Figure S5) were generated using AlphaFold2-multimer(V3) [34, 35] within the colabfold-batch implementation 1.5.5[74]. We used five recycles and five ensembles to produce five models for each sequence. The model with the best ranking after relaxation was selected for further analysis. Templates were not used in this process.

### Conformational sampling with AF2-multimrer(V3)

We enhanced the conformational sampling of AlphaFold2-multimer(V3) using multiple seeds and the *dropout* option that introduces epistemic uncertainty and leads to a wider exploration of potential structures by the model[75]. We used 256 seeds with 1 recycle and 1 ensemble to generate 5 models per seed for a total of 1280 models per sequence. All models were considered for analysis. We used this approach for ancestral prestin sequences and extant representatives of vertebrate prestin: **placentals** (human, gerbil, dolphin), **monotremes** (platypus), **marsupials** (opossum, wombat) **birds** (chicken, egret, finch, pigeon), **reptiles** (alligator, crocodile, rattlesnake, turtle), **amphibians** (frog) and **fish** (ricefish, tetra, zebrafish)

Further conformational sampling was performed by combining dropout with reduction of the depth of the multiple sequence alignment (MSA) used as input, following previous work showing that MSA subsampling can modulate the strength of the co-evolutionary signal and promote exploration of alternative states[40, 42, 63] Specifically, we varied the parameters controlling the number of clustered (max-seq) and extra (max-extra-seq) sequences, using three combinations (64:128, 128:256 and 256:512) and four recycles. We carried out 96 independent seed runs, generating five models per seed for each MSA configuration, for a total of 480 models per sequence. This procedure was applied to the reconstructed ancestors (ancAmn, ancRep, ancMam, ancPla, ancThe) and to HsPres, providing an extended ensemble for comparative analysis of conformational variability.

#### A. Molecular Dynamics Simulations

Representative structural predictions obtained with AF2-muiltimer(V3) for ancAmn, ancRep, ancMam (two models) and HsPres (two models) were selected to generate all-atom MD simulations systems using CHARMM-GUI membrane builder [76, 77, 78]. The proteins were embedded in a 1-palmitoyl-2-oleoyl-sn-glycero-3-phosphocholine (POPC) bilayer with their position and orientation calculated from the OPM server [79]. The systems were solvated with TIP3P water and ions K^+^ and Cl^−^ were added to the final concentration equivalent to 150 mM of KCl. Each system resulted in a rectangular box of dimensions 180x180x155 Å^3^, containing ∼470K atoms, for mammalian systems (ancMam, HsPres), and 220x220x160^3^, containing ∼740K atoms for nonmammalian systems (ancAmn, ancRep). The protonation states of the ionizable residues were selected considering a pH of 7.5 (residues D, E, K, and R were charged and the histidine residues were neutral). Neutral termini N (ACE) and C (CT3) were used in all systems

All simulations were performed with GROMACS 2023.4[80, 81, 82] using the CHARMM36m force field for proteins [83] and the charmm36 force field for lipids[84]. The systems were minimized and subjected to a protocol of constrained NVT and NPT relaxations as proposed by the CHARMM-GUI membrane builder tool. Finally, NPT unconstrained runs of 1.2 μs were performed. In all cases, the first 200 ns were considered as equilibration and the remaining simulation time as production run. The Berendsen thermostat was used for the constrained relaxation runs, and the v-rescale thermostat was used during the equilibration and production run. In all cases, the temperature was 310 K. For the NPT simulations the Berendsen barostat was used during relaxations, and the c-rescale barostat was used in unconstrained equilibration and production runs. In all cases a semiisotropic barostat with a target pressure of 1 atm was used. In all simulations, the verlet cutoff scheme and a cutoff of 12 Å with a switching function starting at 10 Å, were used for nonbonded interactions along with periodic boundary conditions. A uniform timstep of 2fs was used. The Particle Mesh Ewald method was used to compute long-range electrostatic forces [85]. Hydrogen atoms were constrained using the LINCS algorithm [86].

#### B. Data analysis and visualization

Structural analyses of the predicted models and MD trajectories were performed using VMD[87] together with custom Tcl and Python scripts. Intramonomer distances are reported as combined measurements of the monomers of each dimer, while intermonomer distances correspond to the averaged values between pairs AB and BA.

For the simulations, the first 200 ns were considered equilibration, and the remaining time was treated as the production phase used for data analysis. Secondary structure assignments, including monitoring of Cα2 conformations, were obtained using DSSP[88]. [88]. Molecular renderings were performed with VMD and ChimeraX[89].

##### 1. Contacts maps

Contact maps were generated from the MD trajectories using CONAN[90] considering all the heavy atoms of the protein and a cut-off point of 10 Å. For intramonomer contacts, the maps of both monomers were averaged into one map. For intermonomer contacts, the A-B and B-A maps were averaged into one map.

##### 2. Lipid occupancy in the intermonomer region

The occupancy of lipids inside the intermonomer region was recorded as a time series where *O*(*t*) ≥ 1 if there are phosphorus atoms of POPC molecules located within 10 Å of the Cα atoms of residues 489 to 504 (residue numbering based on HsPres but adapted to equivalent residues in nonmammalians) of both chains.

##### 3. Cl^−^ Occupancy and residence times

The residence times of Cl^−^ ions within each binding site were calculated using occupancy data from our MD simulations. The occupancy was recorded as a time series, where *O*(*t*) ≥ 1 if Cl^−^ ions were located within 7 Å of the Cα atoms of residues 97, 101, 136, 137, 397, 398 and 399 (residue numbering based on HsPres but adapted to equivalent residues in nonmammalians) of each monomer at time *t*, and *O*(*t*) = 0 if absent. A residence event was defined as a continuous sequence where *O*(*t*) ≥ 1. The residence time τ for each event was calculated as:

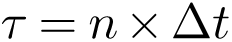

where *n* is the number of consecutive frames with *O*(*t*) ≥ 1, and Δ*t* is the time between frames (20 ps).

## DATA AVAILABILITY

All the outputs of the ASR and AF2-multimerV3 structure predictions and MD trajectories can be found at DOI:10.5281/zenodo.15197370

## AUTHOR CONTRIBUTIONS

**Nicolas Fuentes-Ugarte**: Investigation, Formal Analysis, Visualization, Writing – Review & Editing. **Tiaren Ruiz-Rojas**: Investigation, Formal Analysis, Visualization, Writing – Review & Editing. **Felipe Garcia-Olave: Formal Analysis, Writing – Review & Editing**. **Alvaro Ruiz-Fernandez**: Writing – Review & Editing. **Jose Antonio Garate**: Writing – Review & Editing. **Victor Castro-Fernandez**: Investigation, Conceptualization, Supervision, Writing – Original Draft Preparation, Writing – Review & Editing. **Raul Araya-Secchi**: Investigation, Conceptualization, Funding Acquisition, Writing – Original Draft Preparation, Writing – Review & Editing, Project Administration.

## ACKNOWLEDGMENTS

This work is supported by the following grants: ANID FONDECYT Regular 1231164 and ANID PAI 77200112 (RAS),ANID FONDECYT Regular 1221260 (JAG), ANID FONDECYT De Iniciación 11221268 (ARF), ANID FONDECYT Regular 1221667 (VCF), ANID Beca Doctorado Nacional 21221449 (NFU) and Financiamiento Basal para Centros Cientificos y Tecnologicos de Excelencia de ANID” Centro Ciencia& Vida, FB210008 (to Fundación Ciencia & Vida). The authors thank Felipe Engelberger for his helpful advice and discussion.

