## Supplementary Material for "From Transporter to Motor: Evolutionary and Structural Insights into the Emergence of Prestin’s Area-Motor Activity in Mammals"


**Nicolas Fuentes-Ugarte***

*Departamento de Biologia, Facultad de Ciencias, Universidad de Chile, Santiago,Chile.*

**Tiaren Ruiz-Rojas***

*Facultad de Ingenieria, Tecnologia y Diseño, Universidad San Sebastian, Santiago, Chile*

**Felipe Garcia-Olave**

*Programa de doctorado en Biologia Computacional. Universidad San Sebastian, Santiago, Chile.*

**Alvaro Ruiz-Fernandez**

*Computational Biology Lab, Centro Cientifico y Tecnologico de Excelencia, Fundacion Ciencia & Vida, Santiago, Chile, Santiago, Chile.*

**Jose Antonio Garate**

*Facultad de Ingenieria, Tecnologia y Diseño, Universidad San Sebastian, Santiago, Chile*

**Victor Castro-Fernandez**†

*Departamento de Biologia, Facultad de Ciencias, Universidad de Chile, Santiago,Chile.*

**Raul Araya-Secchi**‡

*Facultad de Ingenieria, Tecnologia y Diseño, Universidad San Sebastian, Santiago, Chile*

*Centro Cientifico y Tecnologico de Excelencia, Fundacion Ciencia & Vida, Santiago, Chile*

*These authors contributed equally to this work

### SUPPLEMENTARY DISCUSSION

#### Mapping the substitutions to the STAS domain.

The cytoplasmic STAS domain is a structural hallmark of the SLC26A family, and since the publication of the first structure[57] it has been considered an important factor for the movement of the TMD during functional conformational change. This domain is structurally conserved between members of the prestin and SLC26 family and constitutes $\sim 15\%$ of the total buried interface between prestin protomers through STAS–STAS interactions, while forming extensive contacts with the opposing N-terminal domain, reinforcing the importance of STAS in dimer stabilization. It has also been suggested to act as a platform for binding of regulatory proteins such as CFTR and calmodulin[58, 60, 61].

In the STAS domain, the substitutions are broadly distributed as shown in Supplementary Figure S11B. However, focusing only on non-conservative substitutions reveals some clustering (Figure S11CF). Some substitutions can be assigned to regions of interaction between adjacent STAS domains, while others could affect the interaction between the STAS domain and its own or adjacent TMD.

However, the lack of experimental mutational studies or systematic comparisons between the STAS domain of mammalian and nonmammalian prestin or among SLC26A paralogs limits the interpretation of our findings. However, previous domain-swap experiments have shown that replacing the N and C terminals of rat prestin with those of zebrafish prestin resulted in a modest depolarizing shift in the voltage of half-activation ($V_{1/2}$), without affecting the slope factor ($\alpha$)[20]. This observation supports the idea that the STAS domain can modulate the electromechanical response. The clustering of non-conservative substitutions at the STAS–STAS interface and near contacts with the TMD suggests that evolutionary remodeling of this domain may have contributed to tuning the voltage sensitivity and dimer stability in the mammalian lineage.

### SUPPLEMENTARY FIGURES

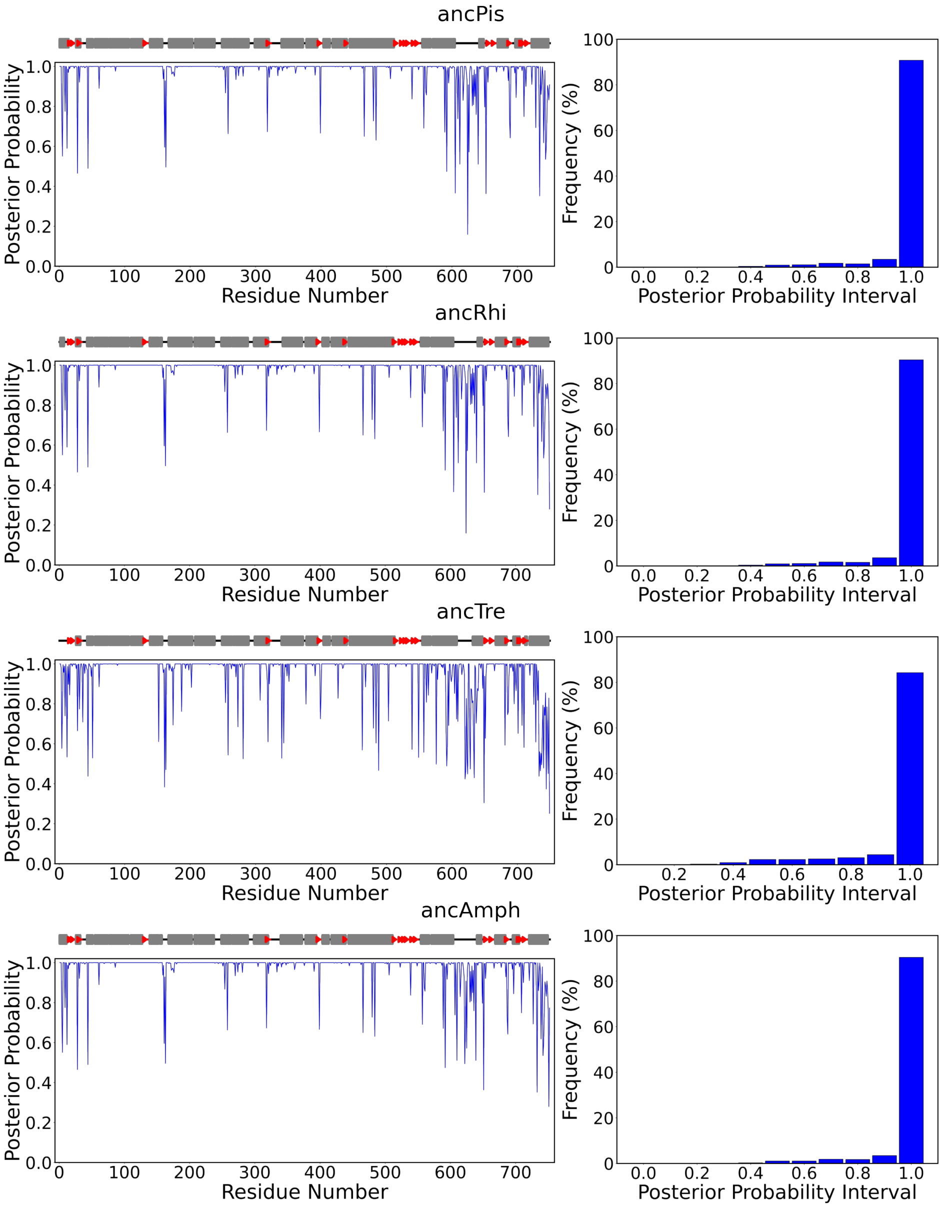


Figure S1. **Robustness of Ancestral Sequence Reconstruction.** The posterior probability for each sequence position (residue number) is shown on the left, the ancestor topology predicted by AlphaFold2 is displayed at the upper left, and the posterior probability distribution for each ancestor is shown on the right. Secondary structure elements are represented as gray tubes for $\alpha$-helices and red arrows for $\beta$-sheets. Both the mean and median posterior probabilities exceed 0.95.

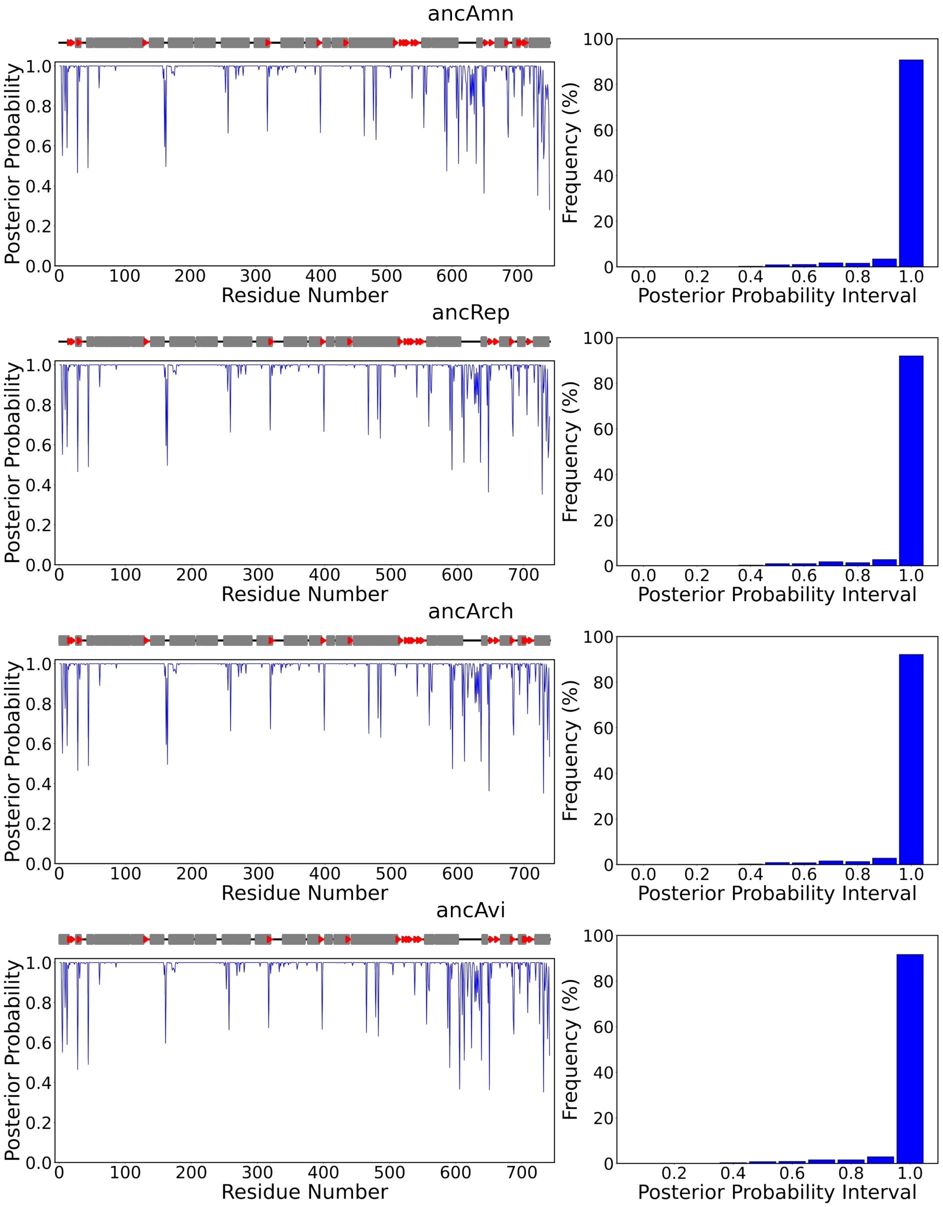


**Continuation of Figure S1.**

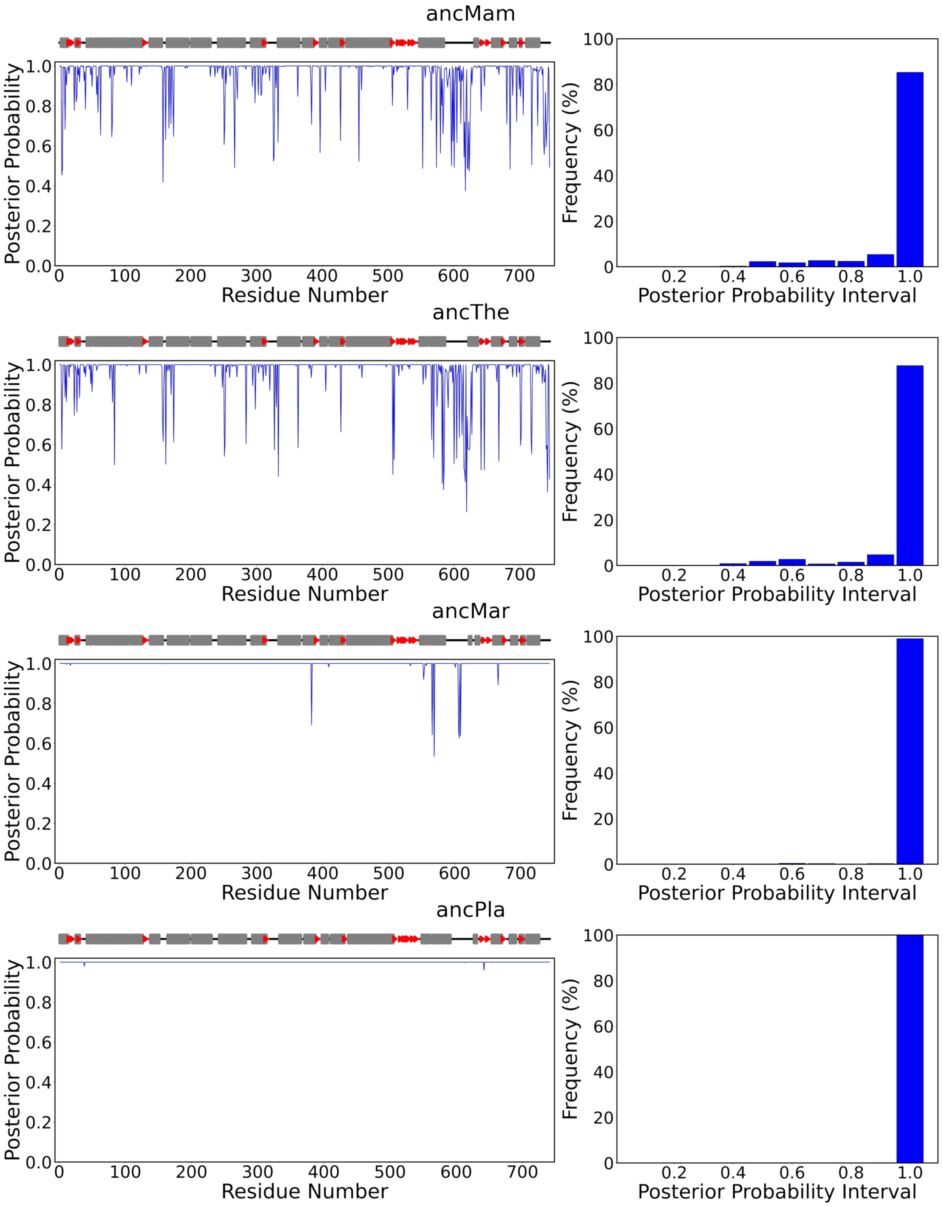


**Continuation of Figure S1.**

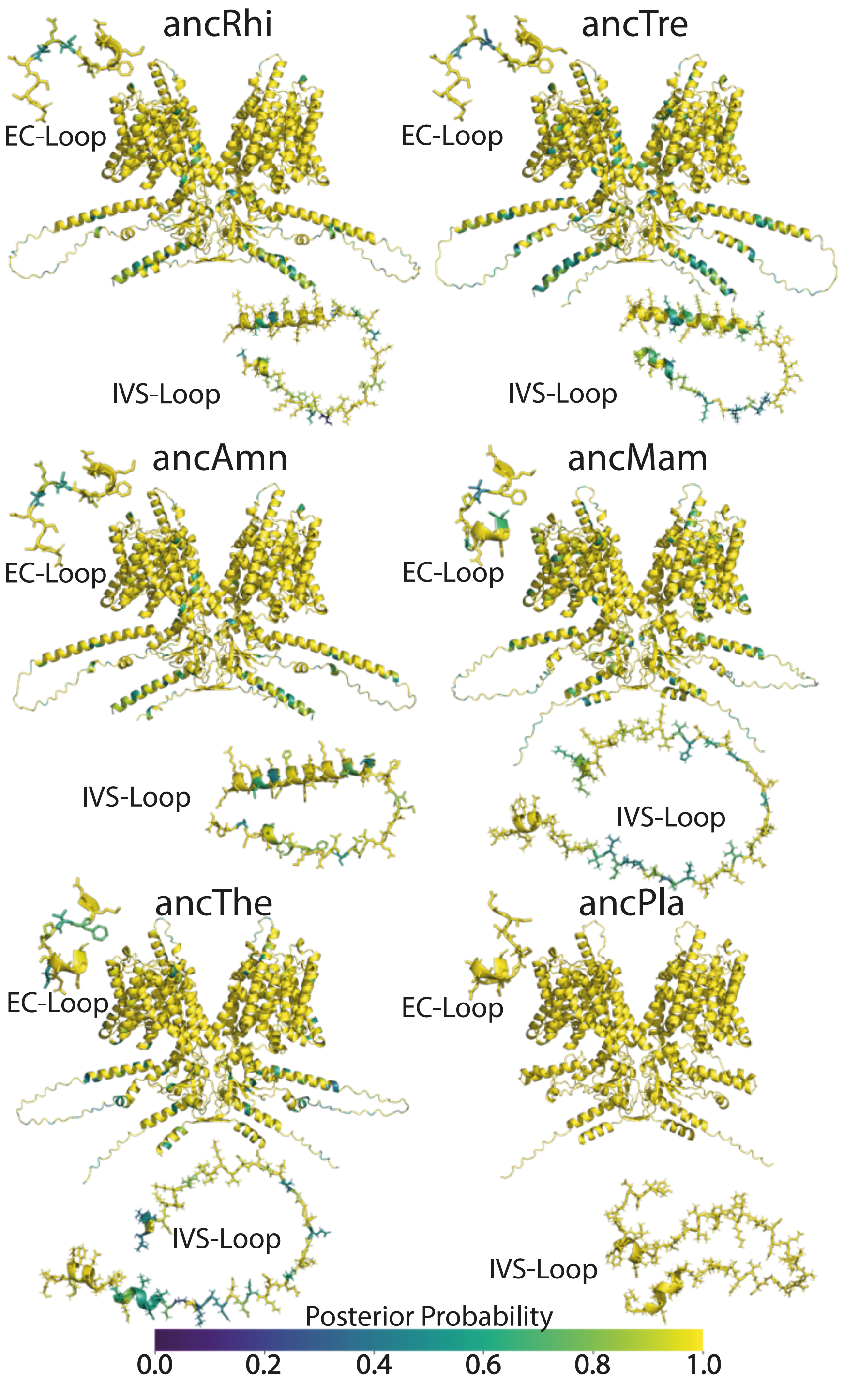


Figure S2. **Analysis of ancestral sequence reconstruction.** The posterior probabilities for each ancestral sequence are mapped to the structural models predicted by AF2-multimer(V3) and colored according to the viridis spectrum. The EC-loop and IVS-loop regions are highlighted in detail to illustrate local confidence and structural consistency across ancestors. A color scale indicating posterior probability values is shown at the bottom of the figure.

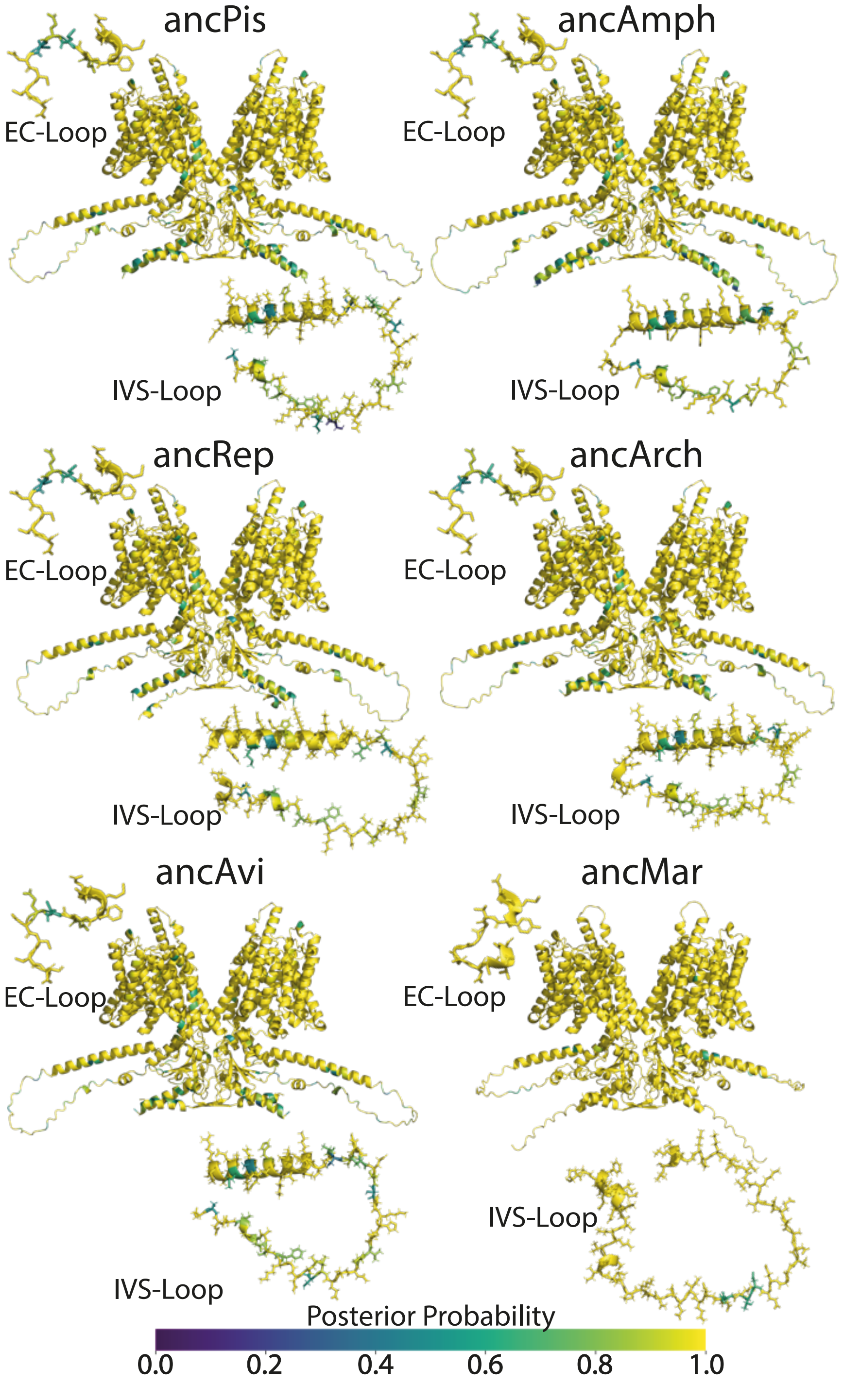


**Continuation of Figure S2.**

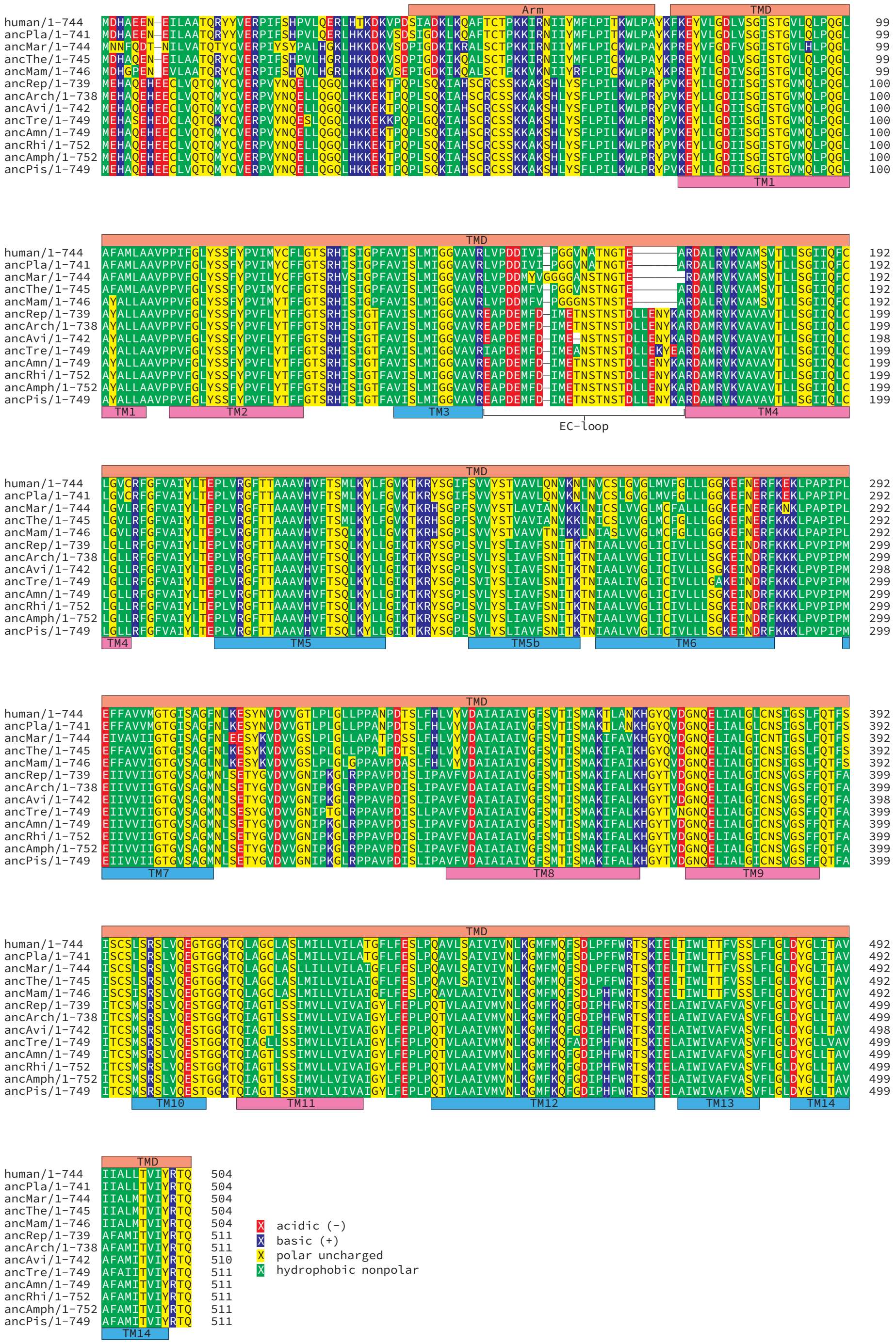


Figure S3. **Ancestral prestin sequences - TMD.** Sequence alignment of the transmembrane domain (TMD) from inferred ancestral prestin sequences, along with human prestin (HsPres). Transmembrane (TM) helices are shown as boxes below the alignment. Helices from the transport domain (core) are depicted in magenta, and those from the scaffold domain (gate) are shown in light blue. The extracellular (EC) loop connecting TM3 and TM4 is indicated by a curly bracket. The positions of the TM helices and the EC-loop are based on the 7LGU human prestin structure. Residues are color-coded according to their hydropathy.

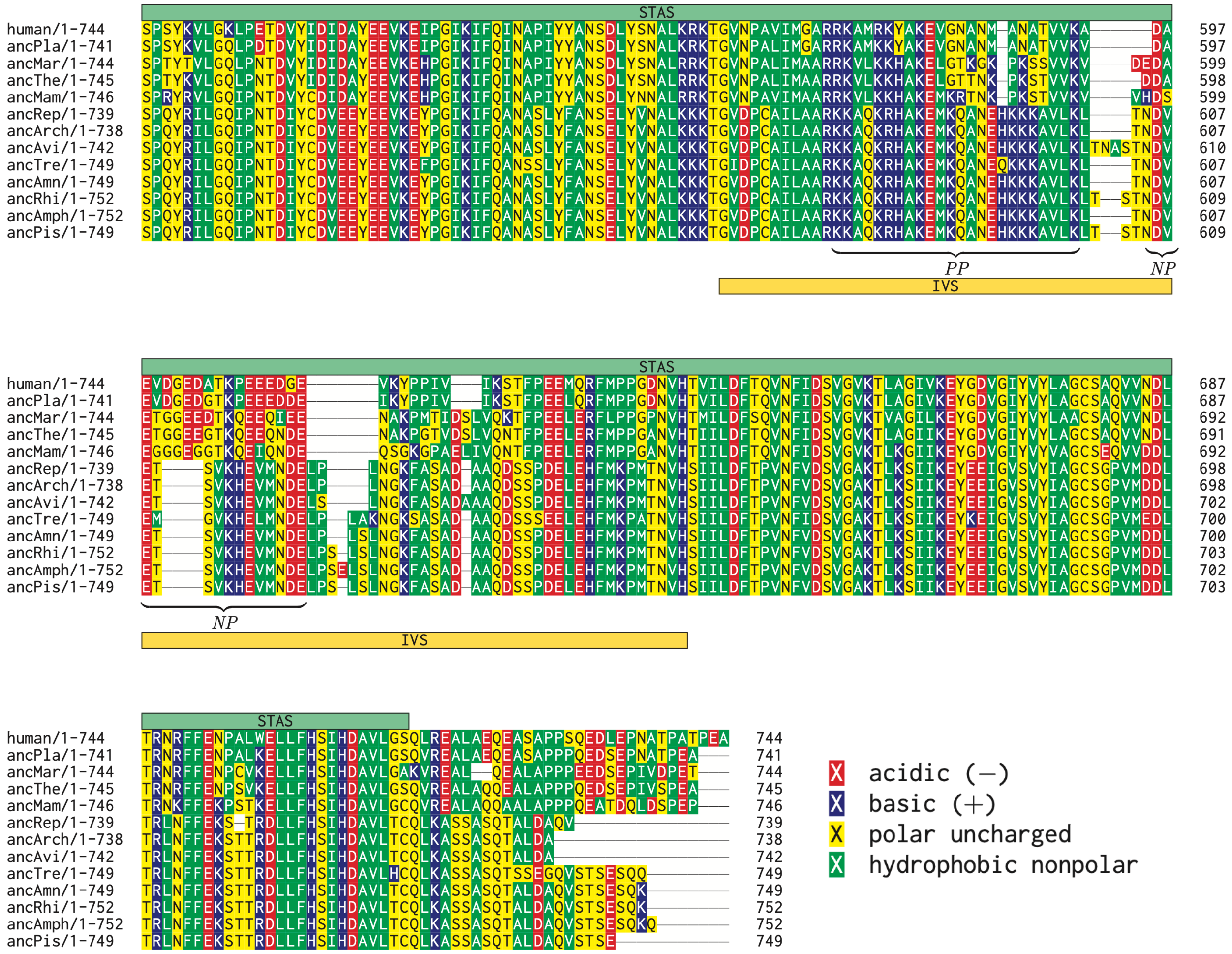


Figure S4. **Ancestral prestin sequences - STAS** Sequence alignment of the STAS domain from inferred ancestral prestin sequences, along with human prestin (HsPres). The IVS is represented as a golden box below the alignment. The positive patch (PP) and the negative patch (NP) are indicated by curly brackets. Residues are color-coded according to their hydropathy.

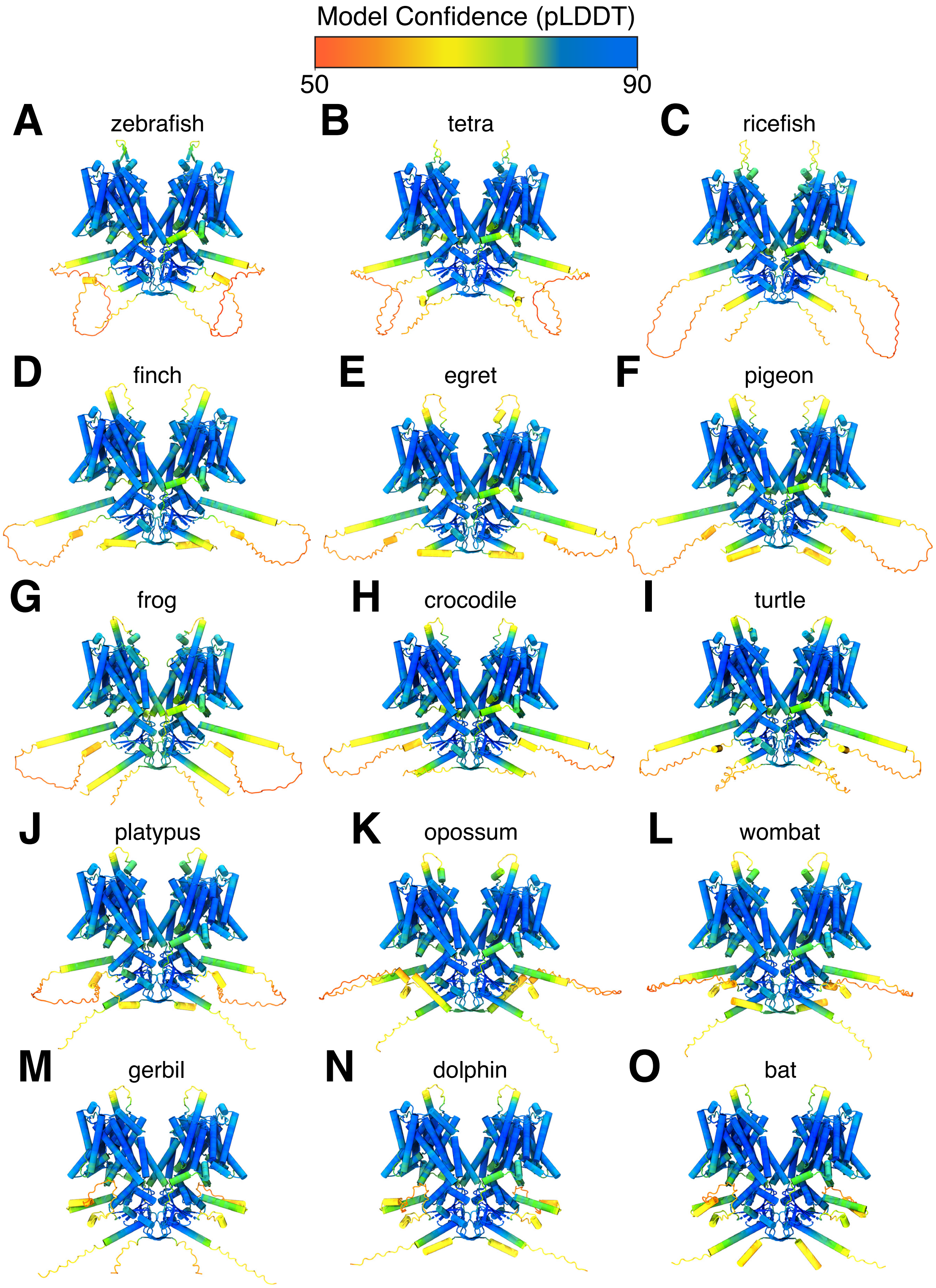


Figure S5. **Reconstruction of nonmammalian and mammalian representative prestin structures using AF2-multimer(V3)**. Models shown in cartoon representation and colored by model confidence (pLDDT).

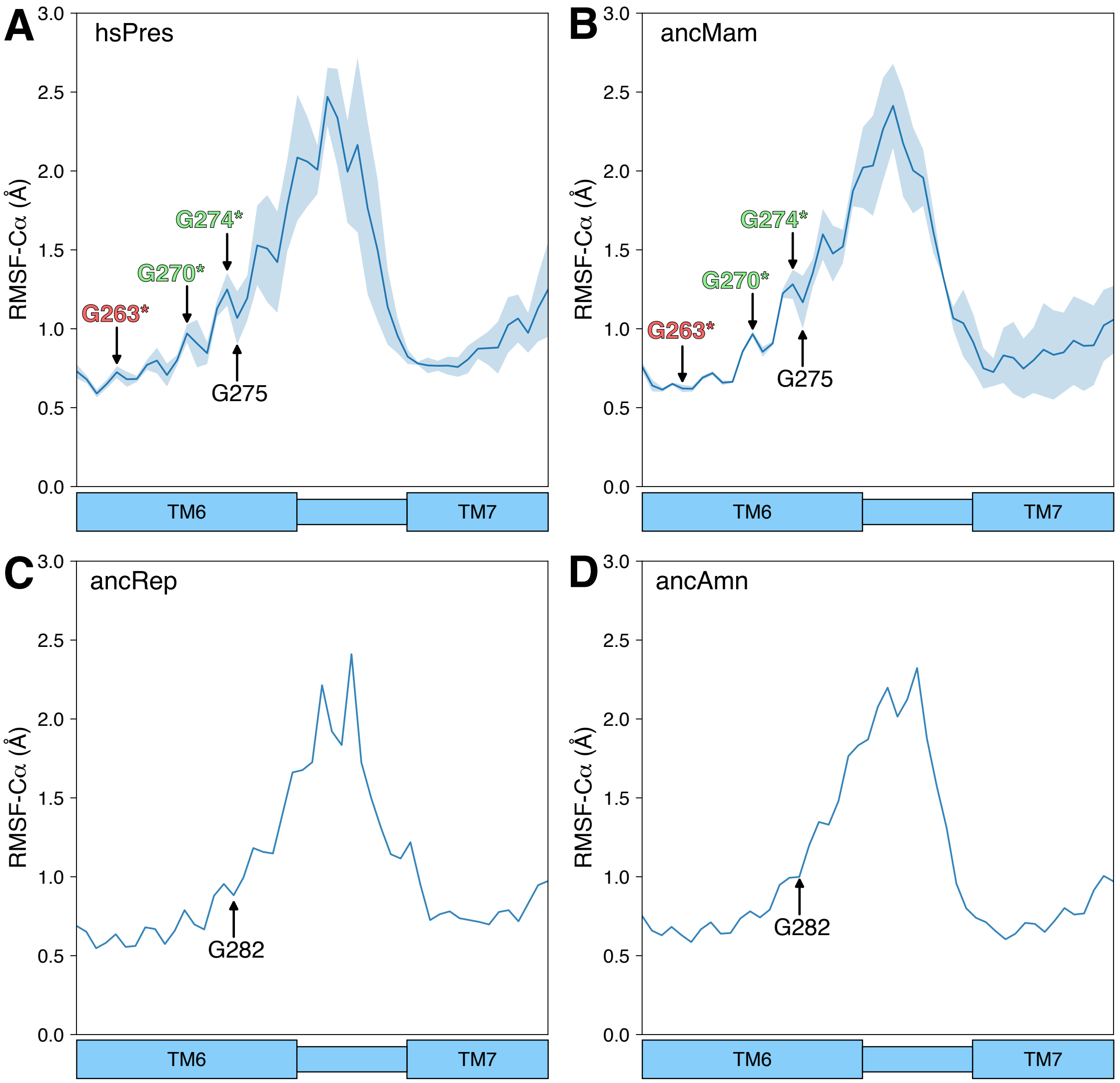


Figure S6. **Structural effect of substitutions in the EC-loop.** RMSF-C$\alpha$ of the EC-loop obtained from the MD simulations of HsPres (A),ancMam (B), ancRep (C) and ancAmn (D). In panels A and B, the key substitutions for glycine are indicated by arrows and colored as in Figure 1.

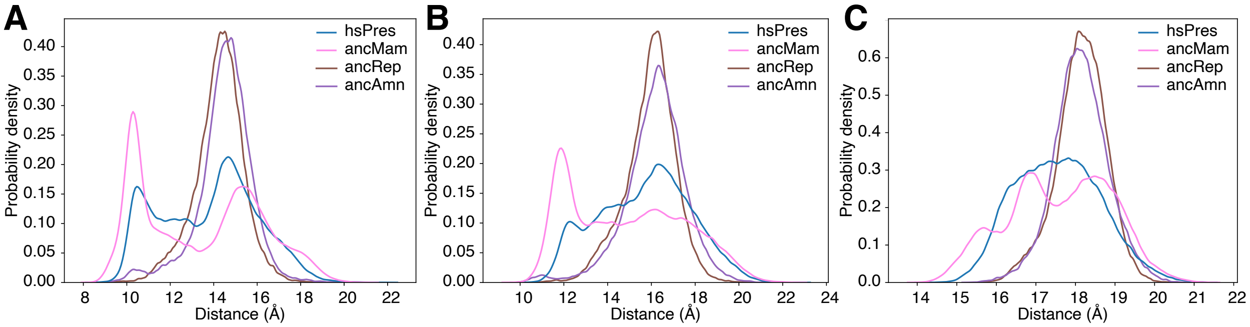


Figure S7. **Structural effect of substitutions in TM6-TM12.** (A) Distance between the C$\alpha$ atoms of residues D277 (TM6) and K449 (TM12). (B) Distance between the C$\alpha$ atoms of residues E280 (TM6) and K449 (TM12) (numbering based on HsPres but measured for the equivalent residues in ancAmn and ancRep) and (C) distance between the center of mass of TM6 and TM12.

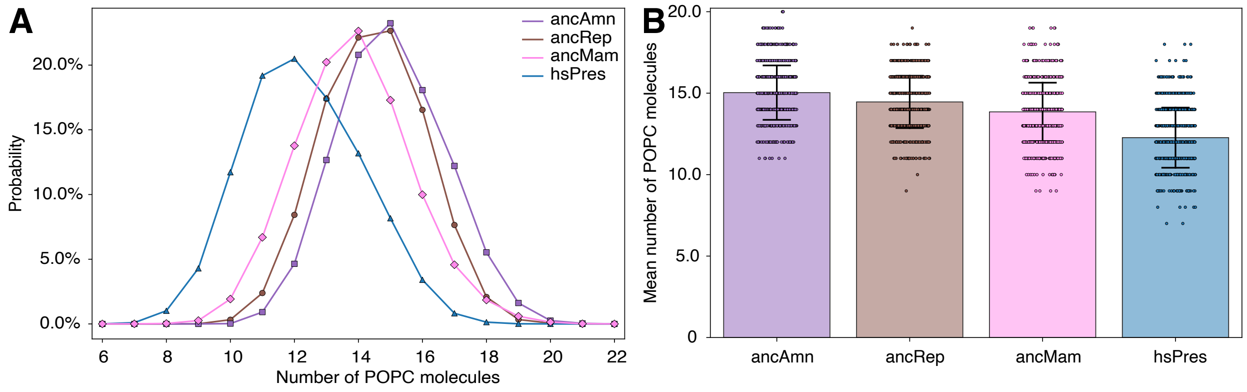


Figure S8. **Lipid occupancy in the intermonomer space.** (A) Distribution of lipid occupancies in the intermonomer space obtained from MD simulations. (B) Mean lipid occupancies (bars ± SD) with individual simulation values shown as circles (see methods for details).

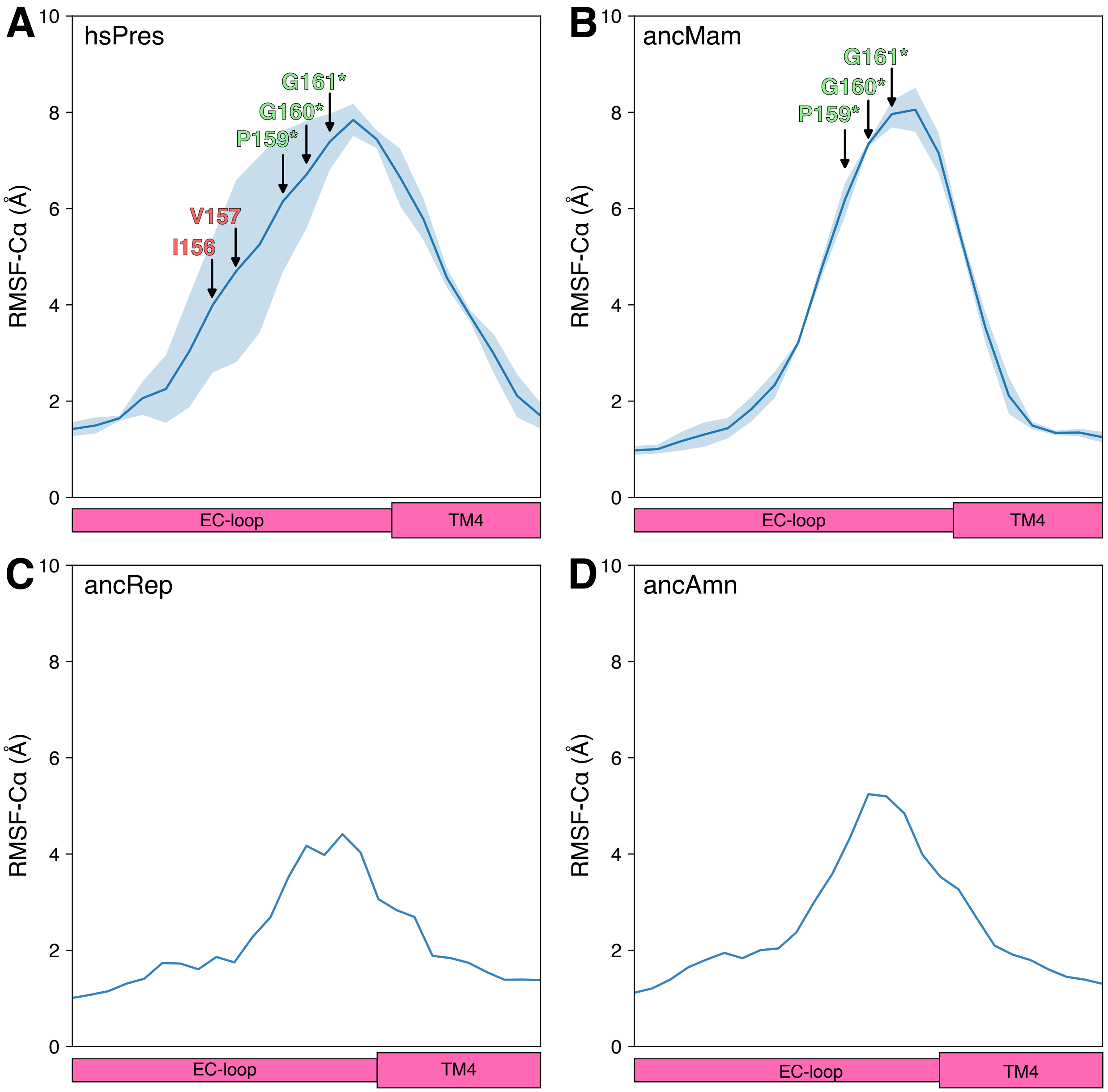


Figure S9. **Structural effect of substitutions in the EC-loop.** RMSF-C$\alpha$ of the EC-loop obtained from the MD simulations of HsPres (A),ancMam (B), ancRep (C) and ancAmn (D). In panels A and B the substitutions that give rise to the PGG motif are indicated by arrows and colored as in Figure 1.

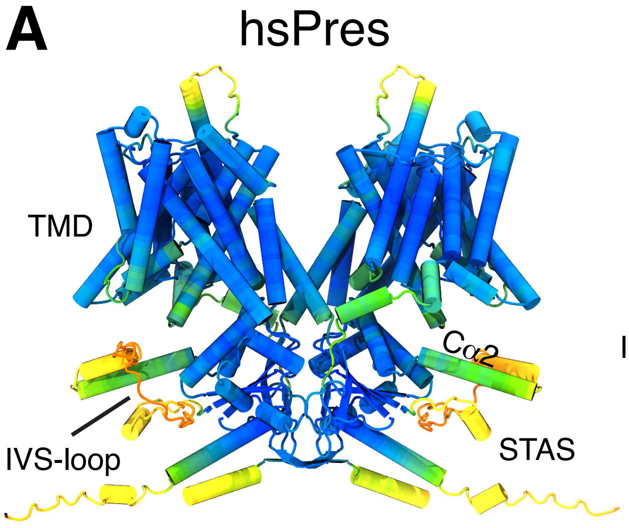


Figure S10. **Comparison between HsPres and SLC26A3.** (A) HsPres model presented and colored as in Figure 2. (B) Rendering of SLC26A3 (PDB-ID: 7XUH) shown in cartoon representation with one chain colored gray and the other ochre. In both chains, the IVS-loop is colored orange.

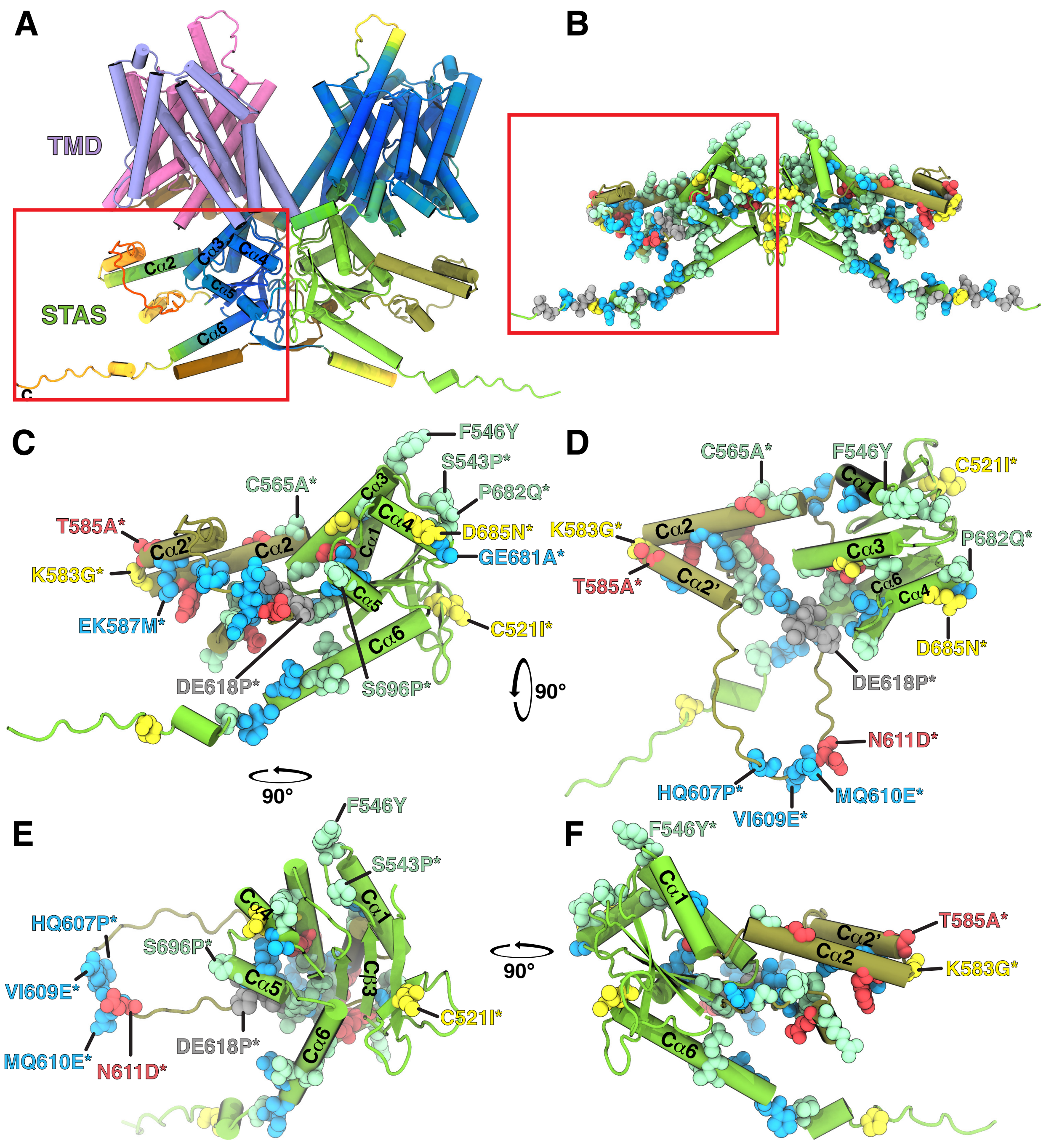


Figure S11. **[Corrected numbering: was Figure 16, now Figure S14.] Substitutions STAS domain** (A)All substitutions found on the STAS domain mapped into the HsPres model. (B-E) Detail of the STAS domain showing only the non-conservative substitutions. (B) Same orientation as in A showing only one monomer. (C) 90° rotation of B along the x-axis to show a top wiew (from the extracellular side). (D) 90° rotation of B along the z-axis. (E) 90° rotation of D along the z-axis. In all panels, the protein is shown in cartoon representation, and the substituted residues are shown in VdW representation. The same color scheme as in Figure 1 (B-D) was used.

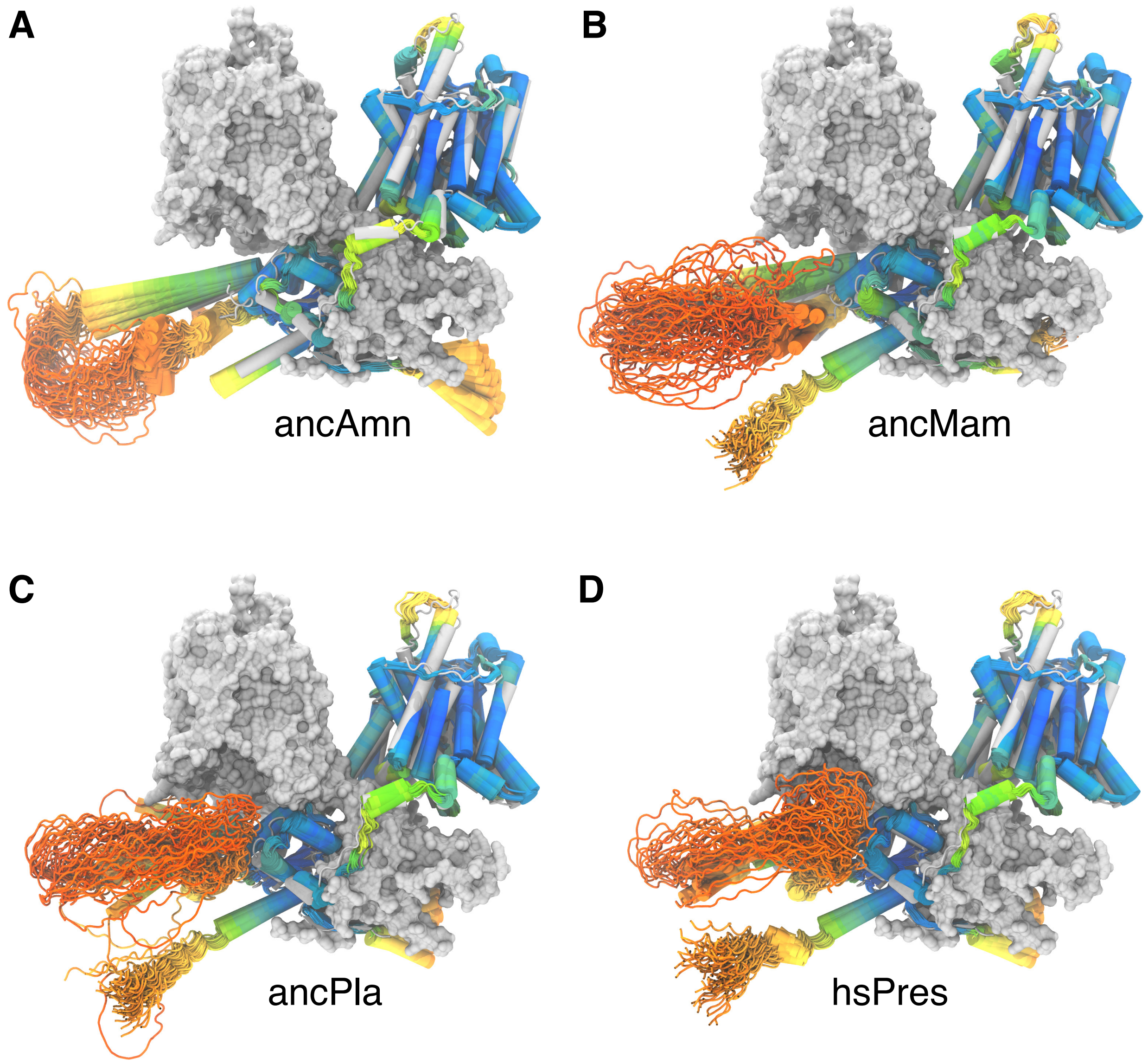


Figure S12. **Conformational sampling of prestin models**(A-D) Representative conformations obtained for human and ancestral prestin models. Only one monomer of the models is shown in cartoon representation colored as in Figure 2. The other monomer corresponds to HsPres structure 7LGU[27] shown as a gray surface.

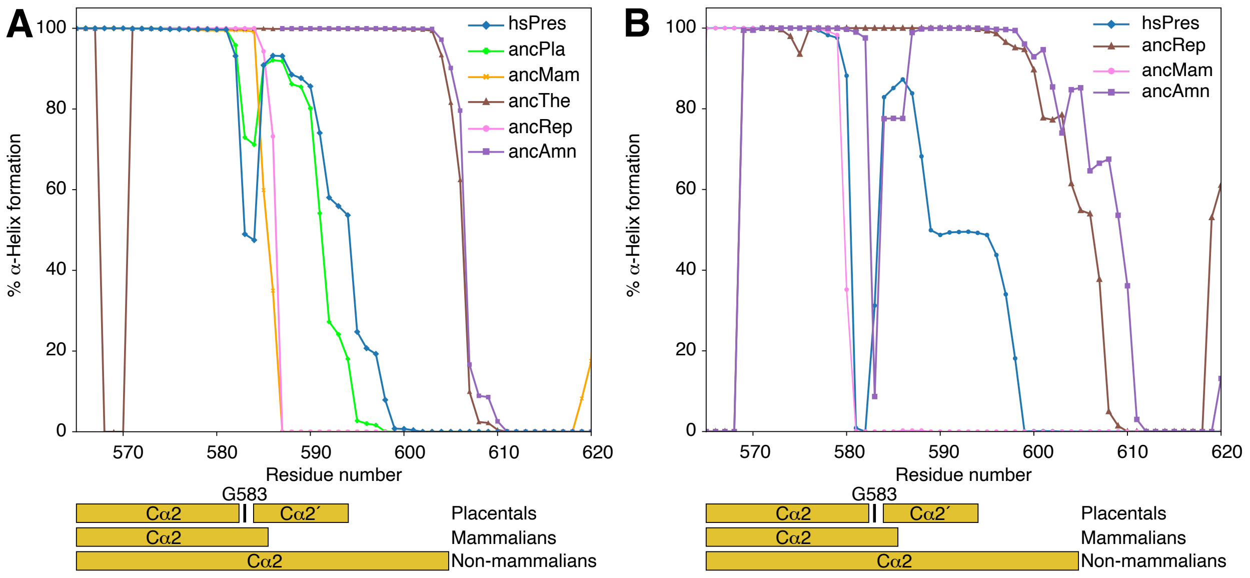


Figure S13. **Conformational dynamics of the C$\alpha$2 helix** (A-B) Percentage of alpha-helix formation for each residue in the IVS of the AF2-multimerV3 models (A) and in the MD simulations (B). The plots show the percentage of models or simulation frames in which each residue is assigned an alpha-helix conformation (H) by DSSP. The dark yellow blocks below each panel highlight the length of the helix for each prestin group (Placental, Mammalians and nonmammalians) and the location of residue G583. The residue numbers on both panels correspond to HsPres.

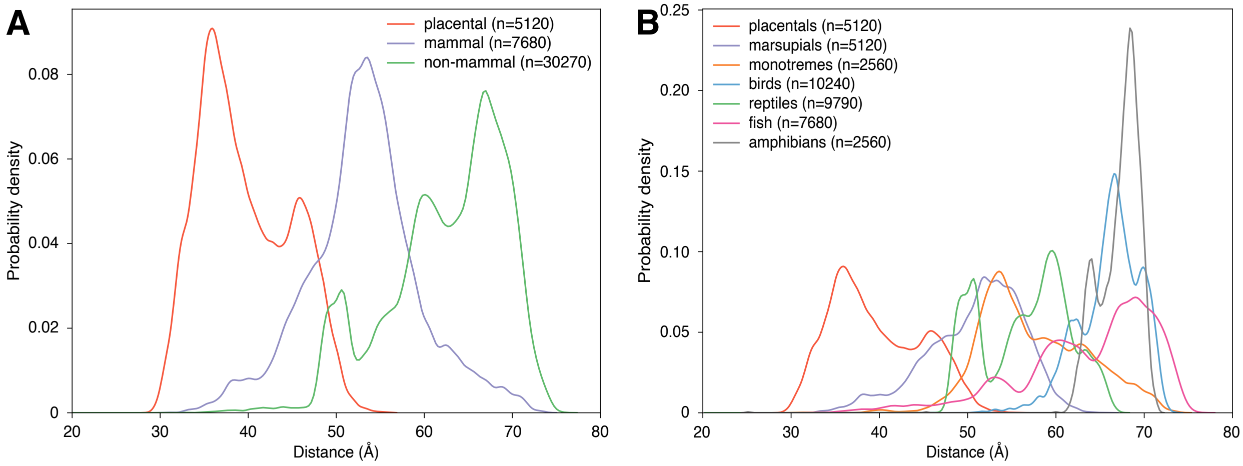


Figure S14. **Distribution of the intermonomer IVS − Cl B.S. distance across vertebrates** (A-B) Distribution of the distance between the center of mass of the IVS-loop of one monomer and the chloride binding site of the adjacent monomer obtained from models predicted with AF2-multimer(V3) across vertebrate clades. (A) Broad grouping into placental mammals, other mammals, and nonmammalian species. (B) Same data set resolved by the major vertebrate lineages: placentals, marsupials, monotremes, birds, reptiles, fish, and amphibians. The parentheses in the legend indicate the number of models per group or clade (see methods for details).

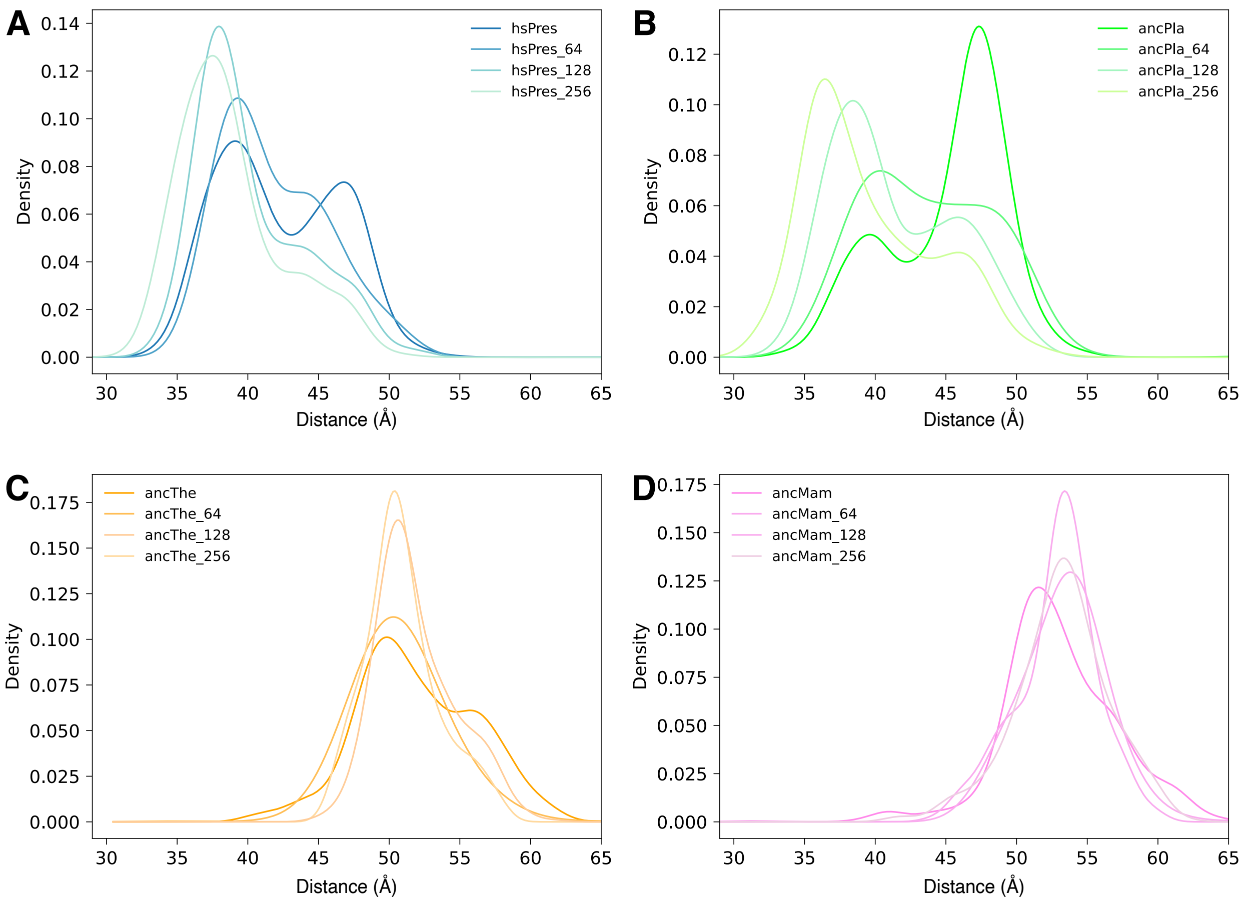


Figure S15. **Effect of MSA depth on the distribution of the intermonomer IVS − Cl B.S. distance.** (A-B) Distribution of the distance between the center of mass of the IVS-loop of one monomer and the chloride binding site of the adjacent monomer obtained from models predicted with AF2-multimer(V3) with for HsPres (A), ancPla (B), ancThe (C) and ancMam (D) (see methods for details).

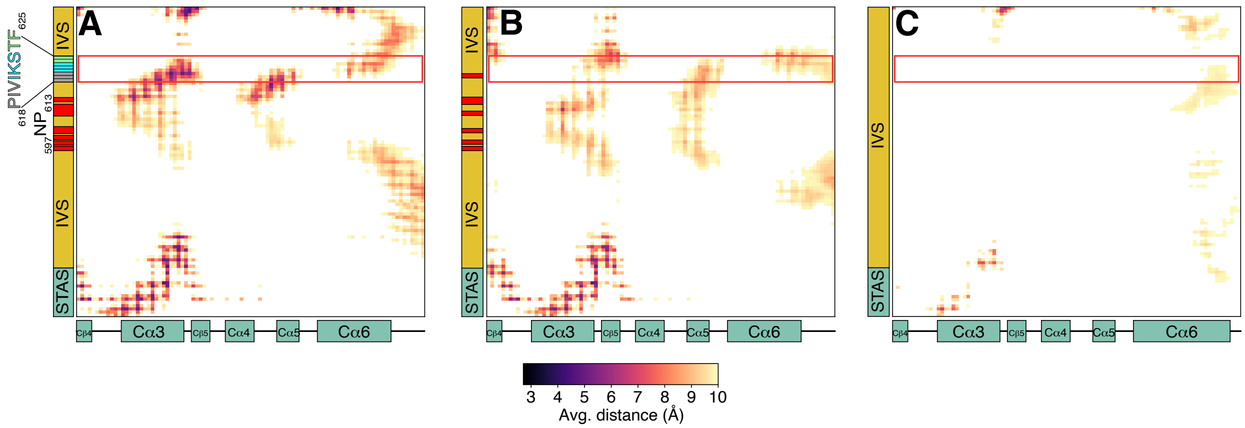


Figure S16. **Effect of substitutions on the interactions between the IVS and the rest of the STAS domain** (A-C) Average distance maps between the IVS and the rest of the STAS domain obtained from the MD simulations of HsPres (A), ancMam (B) and ancAmn (C). Prestin topology features are colored as in Figure 1

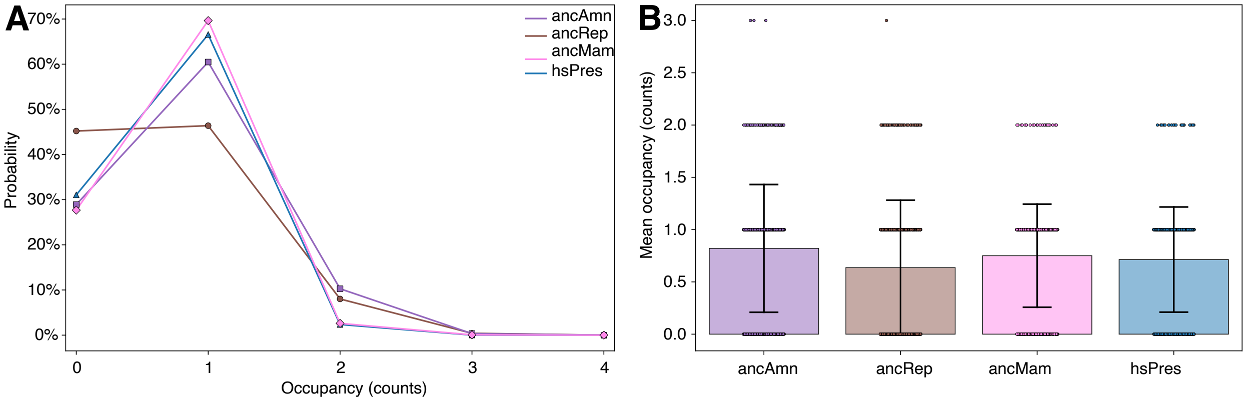


Figure S17. **Chloride occupancy in $Cl^-$ binding site.** (A) Distribution of $Cl^-$ occupancies in the $Cl^-$ binding site obtained from MD simulations. (B) Mean $Cl^-$ occupancies (bars ± SD) with individual simulation values shown as circles (see methods for details).

### SUPPLEMENTARY TABLES

Table S1. Sequence identity between the inferred ancestral prestin sequences sequences and HsPres.

| | ancPis | ancRhi | ancTre | ancAmph | ancAmn | ancRep | ancArch | ancAvi | ancMam | ancThe | ancMar | ancPla | HsPres |
|---|---|---|---|---|---|---|---|---|---|---|---|---|---|
| ancPis | - | 99.60% | 92.70% | 98.80% | 99.10% | 98.50% | 98.40% | 98.00% | 66.90% | 63.90% | 61.20% | 60.60% | 59.80% |
| ancRhi | | - | 93.00% | 99.20% | 99.50% | 98.10% | 98.00% | 97.60% | 67.10% | 63.90% | 61.20% | 60.60% | 59.80% |
| ancTre | | | - | 93.10% | 93.50% | 92.40% | 92.30% | 91.60% | 66.70% | 63.90% | 61.30% | 61.00% | 60.00% |
| ancAmph | | | | - | 99.60% | 98.30% | 98.10% | 97.20% | 67.00% | 63.80% | 61.10% | 60.50% | 59.80% |
| ancAmn | | | | | - | 98.70% | 98.50% | 97.60% | 67.20% | 64.00% | 61.30% | 60.70% | 60.00% |
| ancRep | | | | | | - | 99.60% | 98.70% | 66.90% | 64.00% | 61.40% | 60.70% | 60.10% |
| ancArch | | | | | | | - | 99.10% | 66.90% | 64.00% | 61.40% | 60.70% | 60.10% |
| ancAvi | | | | | | | | - | 66.90% | 63.80% | 61.20% | 60.60% | 59.90% |
| ancMam | | | | | | | | | - | 90.40% | 82.10% | 82.60% | 80.00% |
| ancThe | | | | | | | | | | - | 89.00% | 89.10% | 86.10% |
| ancMar | | | | | | | | | | | - | 81.20% | 79.00% |
| ancPla | | | | | | | | | | | | - | 96.00% |
| HsPres | | | | | | | | | | | | | - |

Table S2. Average posterior probabilities of studied Prestin segments.

| Ancestor | ECloop | IVS loop | TM5b | TM6 | TM7 | TM8 | TM11 | TM12 | TM13 | TM14 |
|---|---|---|---|---|---|---|---|---|---|---|
| ancPis | 0.918 | 0.902 | 0.969 | 0.992 | 0.983 | 0.998 | 1.000 | 0.987 | 0.962 | 0.996 |
| ancRhi | 0.918 | 0.902 | 0.969 | 0.992 | 0.981 | 0.998 | 1.000 | 0.987 | 0.962 | 0.996 |
| ancTre | 0.902 | 0.856 | 0.956 | 0.961 | 0.968 | 0.995 | 0.990 | 0.981 | 0.926 | 0.985 |
| ancAmn | 0.918 | 0.925 | 0.969 | 0.992 | 0.983 | 0.998 | 1.000 | 0.987 | 0.962 | 0.996 |
| ancMam | 0.895 | 0.865 | 0.944 | 0.957 | 0.964 | 0.993 | 0.982 | 0.978 | 0.999 | 0.998 |
| ancThe | 0.899 | 0.839 | 0.936 | 0.993 | 0.977 | 0.982 | 0.984 | 0.997 | 1.000 | 0.999 |
| ancPla | 1.000 | 1.000 | 1.000 | 1.000 | 1.000 | 1.000 | 1.000 | 1.000 | 1.000 | 1.000 |

Table S3. **Substitutions found in N-terminal domain. Pink higlight** Earliest substitutions in the evolutionary pathway. **Gray highlight:** Three-step substitutions. **Blue highlight:** Two-step substitutions. **Light green highlight:** Early substitutions (from the amniote ancestor to the mammalian ancestor). **Yellow highlight:** Intermediate substitutions (from the mammal ancestor to the therian ancestor). **Light red highlight:** Late substitutions (from the therian ancestor to the placental ancestor). Non-conservative substitutions are marked with an asterisk (*). The estimated time of existence for the ancestral proteins were extracted from TimeTree[91]. Cells containing A/B indicate positions where the most probable ancestral sequences and AltALL sequences differ.

| | HsA6 (Outgroup) | ancPis (415 MYA) | ancRhi (408.5 MYA) | ancTre (351.7 MYA) | ancAmn (318.9 MYA) | ancMam (180.1 MYA) | ancThe (160 MYA) | ancPla (99.2 MYA) | HsA5 (Extant) |
|---|---|---|---|---|---|---|---|---|---|
| N-Terminal | G2 | E2 | E2 | E2 | E2 | D2 | D2 | D2 | D2 |
| | D5 | Q5/S5 (0.551/0.230) | Q5/S5 (0.551/0.230) | S5 | Q5/S5 (0.551/0.23) | P5/Q5 (0.482/0.439) | Q5 | E5 | E5* |
| | S7 | H7 | H7 | H7 | H7 | N7 | N7 | N7 | N7* |
| | G8 | E8 | E8 | E8 | E8 | - | - | - | - |
| | M22 | C10 | C10 | C10 | C10 | V9/I9 (0.683/0.290) | I9 | I9 | I9 |
| | L24 | V12/A12 (0.590/0.366) | V12/A12 (0.590/0.366) | A12/T12 (0.534/0.314) | V12/A12 (0.590/0.366) | A11 | A11 | A11 | A11* |
| | R25 | Q13 | Q13 | Q13 | Q13 | A12 | A12 | A12 | A12* |
| | D28 | M16 | M16 | K16 | M16 | R15 | R15 | R15 | R15* |
| | H30 | C18 | C18 | C18 | C18 | C17 | C17 | Y17 | Y17* |
| | L35 | V23 | V23 | V23 | V23 | I22 | I22 | I22 | I22 |
| | L36 | Y24 | Y24 | Y24 | Y24 | F23/Y23 (0.777/0.223) | F23/Y23 (0.746/0.254) | F23 | F23 |
| | N37 | N25 | N25 | N25 | N25 | S24 | S24 | S24 | S24 |
| Nα1 | Q38 | Q26 | Q26 | Q26 | Q26 | H25 | H25 | H25 | H25 |
| | E39 | E27 | E27 | E27 | E27 | Q26 | P26 | P26 | P26* |
| | H40 | L28 | L28 | S28 | L28 | V27 | V27 | V27 | V27* |
| | E42 | Q30 | Q30 | Q30 | Q30 | H29 | H29 | Q29 | Q29 |
| | E43 | G31 | G31 | G31/E31 (0.774/0.207) | G31 | G30 | G30 | E30 | E30* |
| | L44 | Q32 | Q32 | Q32 | Q32 | R31 | R31 | R31 | R31* |
| Loop Nα1-Nα2 | - | E37 | E37 | E37 | E37 | D36 | D36 | D36 | D36 |
| | - | T39 | T39 | K39 | T39 | V38 | V38 | V38 | V38 |
| | A50 | Q41 | Q41 | Q41 | Q41 | E40 | D40 | D40 | D40 |
| | P51 | P42 | P42 | P42 | P42 | P41 | P41 | S41 | S41* |
| Nα2 | R52 | L43 | L43 | L43 | L43 | I42 | I42 | I42 | I42 |
| | T53 | S44/G44 (0.490/0.485) | S44/G44 (0.490/0.485) | G44/S44 (0.438/0.323) | S44 | G43 | G43 | G43 | A43* |
| | H54 | Q45 | Q45 | Q45 | Q45 | D44 | D44 | D44 | D44* |
| | W56 | I47 | I47 | I47 | I47 | I46 | I46 | L46 | L46 |
| | R57 | A48 | A48 | A48 | A48 | K47 | K47 | K47 | K47* |
| | T58 | H49 | H49 | H49 | H49 | Q48 | Q48 | Q48 | Q48* |
| | W59 | S50 | S50 | S50 | S50 | A49 | A49 | A49 | A49* |
| | L60 | C51 | C51 | C51/F51 (0.529/0.462) | C51 | L50 | L50 | F50 | F50* |
| Loop Nα2-Nα3 | Q61 | R52 | R52 | R52 | R52 | S51 | S51 | T51 | T51* |
| Nα3 | S63 | S54 | S54 | S54 | S54 | T53 | T53 | T53 | T53 |
| | R64 | S55 | S55 | S55 | S55 | P54 | P54 | P54 | P54* |
| | A67 | A58 | A58 | A58 | A58 | V57 | I57 | I57 | I57* |
| | Y68 | K59 | K59 | K59 | K59 | K58 | R58 | R58 | R58 |
| | A69 | S60 | S60 | S60 | S60 | N59/H59 (0.767/0.204) | N59 | N59 | N59* |
| | L70 | H61 | H61 | H61 | H61 | I60 | I60 | I60 | I60* |
| | L71 | L62 | L62 | L62 | L62 | I61 | I61 | I61 | I61 |
| | Q73 | S64 | S64 | S64 | S64 | R63/M63 (0.654/0.213) | M63 | M63 | M63* |
| Nα4 | L78 | L69 | L69 | L69 | L69 | C68 | C68 | T68 | T68* |
| Loop Nα4-TMD1 | R83 | R74 | R74 | R74 | R74 | A73 | A73 | A73 | A73* |
| | P85 | P76 | P76 | P76 | P76 | K75 | K75 | K75 | K75* |

Table S4. **Substitutions found in the transmembrane domain (TMD).** Same legend as Table S3

| | HsA6 (Outgroup) | ancPis (415 MYA) | ancRhi (408.5 MYA) | ancTre (351.7 MYA) | ancAmn (318.9 MYA) | ancMam (180.1 MYA) | ancThe (160 MYA) | ancPla (99.2 MYA) | HsA5 (Extant) |
|---|---|---|---|---|---|---|---|---|---|
| TM1 | V86 | V77 | V77 | V77 | V77 | P76 | P76 | F76 | F76* |
| | L90 | L81 | L81 | L81 | L81 | I80/V80 (0.645/0.338) | V80 | V80 | V80 |
| | L94 | I85 | I85 | I85 | I85 | I84 | L84/F84 (0.498/0.282) | L84 | L84 |
| | L95 | I86 | I86 | I86 | I86 | V85 | V85 | V85 | V85 |
| | L98 | I89 | I89 | L89 | I89 | I88 | I88 | I88 | I88 |
| | M93 | M94 | M94 | M94 | M94 | L93 | L93 | L93 | L93 |
| | Y111 | Y102 | Y102 | Y102 | Y102 | Y101 | F101 | F101 | F101 |
| | L113 | L104 | L104 | L104 | L104 | L103 | M103 | M103 | M103* |
| TM2 | F131 | F122 | F122 | F122 | F122 | I121 | I121 | I121 | I121* |
| | F134 | T125 | T125 | T125 | T125 | T124 | C124 | C124 | C124* |
| TM3 | T146 | T137 | T137 | T137 | T137 | P136 | P136 | P136 | P136* |
| | L161 | E152 | E152 | I152/V152 (0.611/0.276) | E152 | L151 | L151 | L151 | L151* |
| | A162 | A153 | A153 | A153 | A153 | V152 | V152 | V152 | V152* |
| EC-Loop | A165 | E156 | E156 | E156 | E156 | D155 | D155 | D155 | D155 |
| | L166 | M157 | M157 | M157 | M157 | M156 | M156 | I156 | I156* |
| | N167 | F158 | F158 | F158 | F158 | F157 | F157/Y157 (0.717/0.258) | V157 | V157* |
| | S169 | I160 | I160 | I160 | I160 | P159 | P159 | P159 | P159* |
| | M170 | M161 | M161 | M161/V161 (0.384/0.384) | M161 | G160 | G160 | G160 | G160* |
| | | E162 | E162 | E162 | E162 | G161 | G161 | G161 | G161* |
| | I171 | T166 | T166 | T166 | T166 | G162 | V162/A162 (0.501/0.376) | V162 | V162* |
| | E173 | S168 | S168 | S168 | S168 | S164 | S164 | A164 | A164* |
| TM4 | T174 | D170 | D170 | D170 | E169 | E169 | E169 | E169 | E169 |
| | A179 | M181 | M181 | M181 | L174 | L174 | M174 | L174 | L174* |
| | S185 | V187 | V187 | V187 | M180 | M180 | M180 | M180 | M180* |
| | T186 | A188 | A188 | A188 | S181 | S181 | S181 | S181 | S181* |
| | I201 | L203 | L203 | L203 | L196 | L196 | L196 | C196 | C196* |
| TM5 | Q230 | Q232 | Q232 | Q232 | Q232 | M225 | M225 | M225 | M225* |
| | F235 | L237 | L237 | L237 | L237 | F230 | F230 | F230 | F230* |
| TM5b | L246 | L248 | L248 | L248 | L248 | L241 | F241 | F241 | F241* |
| | I249 | L251 | L251 | I251 | L251 | V244 | V244 | V244 | V244 |
| | V252 | L254 | L254 | L254 | L254 | T247 | T247 | T247 | T247* |
| | L253 | I255 | I255 | I255 | I255 | V248 | V248 | V248 | V248 |
| | W257 | S259 | S259 | S259 | S259 | T252/A252 (0.711/0.209) | A252/T252 (0.614/0.296) | Q252 | Q252* |
| | L259 | I261 | I261 | I261 | I261 | I254 | V254 | V254 | V254 |
| | P260 | T262 | T262 | T262 | T262 | K255 | K255 | K255 | K255* |
| TM6 | S262 | T264 | T264 | T264 | T264 | L257 | L257 | L257 | L257* |
| | G265 | A267 | A267 | A267 | A267 | A260 | C260 | C260 | C260* |
| | T266 | A268 | A268 | A268 | A268 | S261 | S261 | S261 | S261* |
| | V268 | V270 | V270 | I270 | V270 | V263 | V263 | G263 | G263* |
| | V272 | I274 | I274 | I274 | I274 | M267/L267 (0.490/0.448) | M267 | M267 | M267* |
| | A273 | C275 | C275 | C275 | C275 | C268 | C268 | V268 | V268* |
| | G274 | I276 | I276 | I276 | I276 | F269 | F269 | F269 | F269* |
| | V275 | V277 | V277 | V277 | V277 | G270 | G270 | G270 | G270* |
| | V279 | S281 | S281 | G281 | S281 | G274 | G274 | G274 | G274* |
| | L283 | I285 | I285 | I285 | I285 | F278 | F278 | F278 | F278* |
| | D285 | D287 | D287 | D287 | D287 | E285 | E285 | E285 | E280 |
| TM7 | G297 | M299 | M299 | M299 | M299 | L292 | L292 | L292 | L292* |
| | L299 | I301 | I301 | I301 | I301 | F294 | F294 | F294 | F294* |
| | L300 | I302 | I302 | I302 | I302 | F295 | F295 | F295 | F295* |
| | T301 | V303 | V303 | V303 | V303 | A296 | A296 | A296 | A296* |
| | G304 | I306 | I306 | I306 | I306 | I299 | I299 | M299 | M299* |
| | I308 | V310 | V310 | V310 | V310 | V303 | I303 | I303 | I303 |
| | M312 | M314 | M314 | M314 | M314 | L307 | F307 | F307 | F307* |
| | K315 | S317 | S317 | S317 | S317 | K310 | K310 | K310 | K310* |
| | R317 | T319 | T319 | S319/T319 (0.610/0.364) | T319 | S312 | S312 | S312 | S312 |
| TM8 | I358 | M360 | M360 | M360 | M360 | V353 | V353 | V353 | V353* |
| | I365 | I367 | I367 | I367 | I367 | I360 | I360 | T360 | T360* |
| | F366 | F368 | F368 | F368 | F368 | F361 | F361 | L361 | L361* |
| | L368 | L370 | L370 | L370 | L370 | I363 | I363/V363 (0.584/0.397) | N363 | N363* |
| TM9 | I389 | V391 | V391 | V391 | V391 | I384 | I384 | I384 | I384 |
| | I392 | F394 | F394 | F394 | F394 | L387 | L387 | L387 | L387* |
| TM10 | M402 | M404 | M404 | M404 | M404 | I397/L397 (0.565/0.310) | L397 | L397 | L397* |
| | S410 | S412 | S412 | S412 | S412 | G405 | G405 | G405 | G405* |
| TM11 | A420 | T422 | T422 | L422 | T422 | C415 | C415 | C415 | C415 |
| | S422 | S424 | S424 | S424 | S424 | A417 | A417 | A417 | A417* |
| | L424 | I426 | I426 | I426 | I426 | L419 | L419 | L419 | L419 |
| | I426 | V428 | V428 | V428 | V428 | I421 | I421 | I421 | I421 |
| | V431 | V433 | V433 | L433 | V433 | L426 | L426 | L426 | L426 |
| | L433 | I435 | I435 | I435 | I435 | I428/T428 (0.626/0.209) | I428/T428 (0.663/0.303/) | T428 | T428* |
| | E435 | Y437 | Y437 | Y437 | Y437 | F430 | F430 | F430 | F430 |
| TM12 | A443 | T445 | T445 | T445 | T445 | A438 | A438 | A438 | A438* |
| | A446 | A448 | A448 | A448 | A448 | A441 | S441 | S441 | S441* |
| | I450 | M452 | M452 | M452 | M452 | I445 | I445 | I445 | I445* |
| | R458 | K460 | K460 | K460 | K460 | M453 | M453 | M453 | M453* |
| | S461 | G463 | G463 | A463/G463 (0.569/0.278) | G463 | S456/T456 (0.522/0.435) | S456 | S456 | S456* |
| TM13 | L475 | A477 | A477 | A477 | A477 | T470 | T470 | T470 | T470* |
| | V479 | V481 | V481 | V481 | V481 | T474 | T474 | T474 | T474* |
| | T480 | A482 | A482 | A482 | A482 | T475 | T475 | T475 | T475* |
| | A483 | A485 | A485 | A485 | A485 | S478 | S478 | S478 | S478* |
| TM14 | V494 | L496 | L496 | L496 | L496 | I489 | I489 | I489 | I489 |
| | I498 | A500 | A500 | A500 | A500 | I493 | I493 | I493 | I493* |
| | F499 | F501 | F501 | F501 | F501 | I494 | I494 | I494 | I494* |
| | L501 | M503 | M503 | I503 | M503 | L496 | L496 | L496 | L496* |

Table S5. **Substitutions found in the STAS domain.** Same legend as Table S3

| | HsA6 (Outgroup) | ancPis (415 MYA) | ancRhi (408.5 MYA) | ancTre (351.7 MYA) | ancAmn (318.9 MYA) | ancMam (180.1 MYA) | ancThe (160 MYA) | ancPla (99.2 MYA) | HsA5 (Extant) |
|---|---|---|---|---|---|---|---|---|---|
| Cβ1 | S514 | R516 | R516 | R516 | R516 | R509 | K509/R509 | K509 | K509 |
| | V515 | I517 | I517 | I517 | I517 | V510 | V510 | V510 | V510 |
| | V519 | I521 | I521 | I521 | I521 | I514 | L514 | L514 | L514 |
| Cβ2 | I524 | I526 | I526 | I526 | I526 | V519 | V519 | V519 | V519 |
| | R526 | C528 | C528 | C528 | C528 | C521 | I521 | I521 | I521* |
| Cβ3 | S545 | A547 | A547 | A547 | A547 | I540 | I540 | I540 | I540* |
| Loop Cβ3-Cα1 | T548 | S550 | S550 | S550 | S550 | P543 | P543 | P543 | P543* |
| | V549 | L551 | L551 | L551 | L551 | I544 | I544 | I544 | I544 |
| | F551 | F553 | F553 | F553 | F553 | Y546 | Y546 | Y546 | Y546 |
| Cα1 | E555 | E557/D557 | E557/D557 | E557/D557 | E557/D557 (0.691/0.309) | D550 | D550 | D550 | D550 |
| | S558 | V560 | V560 | V560 | V560 | N553/S553 | S553 | S553 | S553* |
| Cα2 | D570 | C572 | C572 | C572 | C572 | A565 | A565 | A565 | A565* |
| | F571 | A573 | A573 | A573 | A573 | V566 | L566/V566 | L566 | V566* |
| | I573 | L575 | L575 | L575 | L575 | M568 | M568 | M568 | M568* |
| | S574 | A576 | A576 | A576/T576 (0.498/0.463) | A576 | A569 | A569 | G569 | G569* |
| | K577 | K579 | K579 | K579 | K579 | R572 | R572 | R572 | R572 |
| | L580 | Q582 | Q582 | Q582 | Q582 | L575 | L575 | M575 | M575* |
| | Q583 | H585 | H585 | H585 | H585 | H578 | H578 | Y578 | Y578* |
| IVs Loop | L591 | M589/L589 | M589/L589 | M589 | M589/L589 (0.671/0.321) | M582 | L582/M582 | V582 | V582* |
| | Q592 | K590 | K590 | K590 | K590 | K583 | G583 | G583 | G583* |
| | K593 | Q591 | Q591 | Q591 | Q591 | R584 | T584/N584 | N584 | N584 |
| | E594 | A592 | A592 | A592 | A592 | T585 | T585 | A585 | A585* |
| | K596 | E594 | E594 | E594 | E594 | K587 | K587 | M587 | M587* |
| | L617 | V613 | V613 | V611 | V611 | T607 | T606 | T605 | T605* |
| | D619 | H615 | H615 | H613 | H613 | Q609 | Q608/P608 | P607 | P607* |
| | R621 | V617 | V617 | L615 | V615 | I611/V611 | E610 | E609 | E609* |
| | S622 | M618 | M618 | M616 | M616 | Q612 | Q611 | E610 | E610* |
| | N623 | N619 | N619 | N617 | N617 | N613 | N612 | D611 | D611* |
| | Q647 | D635 | D635 | D632 | D632/A632 (0.850/0.113) | E623/D623 | D622/E622 | P618 | P618* |
| | E648 | A636 | A636 | A633/S633 (0.615/0.256) | A633 | L624/S624 | S623/L623 | I619 | I619* |
| | D649 | A637 | A637 | A634/L634 (0.430/0.204) | A634 | I625 | L624/I624 (0.573/0.201) | V620 | V620* |
| | S650 | - | - | - | - | V626 | V625 | I621 | I621* |
| | K651 | Q638 | Q638 | Q635 | Q635 | Q627 | Q626 | K622 | K622 |
| | A652 | D639 | D639 | D636 | D636 | N628 | N627 | S623 | S623* |
| | P653 | S640/T640 | S640/T640 (0.511/0.476) | S637/T637 (0.703/0.240) | S637/T637 (0.511/0.476) | T629 | T628 | T624 | T624 |
| | D654 | S641 | S641 | S638 | S638 | F630 | F629 | F625 | F625* |
| | K659 | E646 | E646 | E643 | E643 | E635 | E634 | Q630 | Q630* |
| | A660 | H647 | H647 | H644 | H644 | R636 | R635 | R631 | R631 |
| | L663 | K650 | K650 | K647 | K647 | P639 | P638 | P634 | P634* |
| | Q665 | M652/G652 | M652/G652 | A649/V649 (0.305/0.237) | M649/G649 (0.363/0.231) | G641 | G640 | G636 | G636* |
| | P666 | T653 | T653 | T650 | T650 | A642 | A641/P641 | D637 | D637* |
| Cβ4 | S670 | S657 | S657 | S654 | S654 | T646 | T645 | T641 | T641 |
| | L671 | I658 | I658 | I655 | I655 | I647 | I646/V646 | V642 | V645 |
| Cα3 | L686 | A673 | A673 | A670 | A670 | V662 | V661 | V657 | V657* |
| | K690 | K677 | K677 | K674 | K674 | K666 | A665* | A661 | A661* |
| | N691 | S678 | S678 | S675 | S675 | G667 | G666 | G662 | G662* |
| | F693 | I680 | I680 | I677 | I677 | I669 | I668 | V664 | V664 |
| | R697 | E684 | E684 | K681 | E681 | G673 | G672 | G668 | G668* |
| | E698 | E685 | E685 | E682 | E682 | D674 | D673 | D669 | D669 |
| | I699 | I686 | I686 | I683 | I683 | V675 | V674 | V670 | V670 |
| Cβ5 | E702 | S689 | S689 | S686 | S686 | Y678 | Y677 | Y673 | Y673* |
| | M705 | I692 | I692 | I689 | I689 | V681 | L680 | L676 | L676 |
| Cα4 | S710 | G697 | G697 | G694 | G694 | E686/A686 | A685 | A681 | A681* |
| | P711 | P698 | P698 | P695 | P695 | Q687 | Q686 | Q682 | Q682* |
| | V713 | M700 | M700 | M697 | M697 | V689 | V688 | V684 | V684* |
| | S714 | D701 | D701 | E698 | D698 | D690 | N689 | N685 | N685* |
| Cα5 | S725 | S712 | S712 | S709 | S709 | P701 | P700 | P696 | P696* |
| | I726 | T713 | T713 | T710/I710 (0.582/0.389) | T710 | S702 | S701/A701 | A697 | A697* |
| | T727 | T714 | T714 | T711 | T711 | T703 | V702 | L698 | L698* |
| | K729 | D716 | D716 | D713 | D713 | E705 | E704 | E700 | E700 |
| Cα6 | H745 | K732 | K732 | K729 | K729 | R721 | R720 | R716 | R716 |
| | P746 | A733 | A733 | A730 | A730 | E722 | E721 | E717 | E717* |
| | R747 | S734 | S734 | S731 | S731/A731 (0.352/0.308) | A723 | A722 | A718 | A718* |
| | P748 | S735 | S735 | S732 | S732 | L724 | L723 | L719 | L719* |
| | P750 | S737 | S737 | S734/A734 (0.560/0.398) | S734 | Q726 | Q725 | E721 | E721* |
| | S752 | T739 | T739 | T736/A736 (0.465/0.248) | T736 | A728/E728 | E727 | E723 | E723* |

Table S6. **Substitutions found in the C-terminal tail.** Same legend as Table S3

| | HsA6 (Outgroup) | ancPis (415 MYA) | ancRhi (408.5 MYA) | ancTre (351.7 MYA) | ancAmn (318.9 MYA) | ancMam (180.1 MYA) | ancThe (160 MYA) | ancPla (99.2 MYA) | HsA5 (Extant) |
|---|---|---|---|---|---|---|---|---|---|
| C-Terminal | V754 | L741 | L741 | S738 | L738 | L730 | L729 | S725 | S725* |
| | S755 | D742 | D742 | E739/D739 | D739 | A731 | A730 | A726 | A726* |
| | V756 | A743 | A743 | G740 | A740 | P732 | P731 | P727 | P727* |
| | T757 | Q744 | Q744 | Q741 | Q741 | P733 | P732 | P728 | P728* |
| | R758 | V745 | V745 | V742 | V742 | P734 | P733 | P729 | S729* |
| | L759 | S746 | S746 | S743 | S743 | Q735 | Q734 | Q730 | Q730 |
| | - | T747 | T747 | T744/S744 | T744 | E736 | E735 | E731 | E731* |
| | - | S748 | S748 | S745 | S745 | A737/D737 (0.628/0.210) | D736 | D732 | D732* |
| | - | E749 | E749 | E746/D746 (0.644/0.221) | E746 | T738/S738 | S737 | S733 | L733* |
| | - | - | S750 | S747 | S747 | D739/E739 | E738 | E734 | E734* |
| | - | - | Q751/H751 (0.757/0.235) | Q748 | Q748/H748 (0.757/0.235) | Q740 | P739 | P735 | P735* |
| | - | - | K752 | Q749 | K749 | L741 | I740 | N736 | N736* |
| | - | - | - | - | - | D742 | V741/A741(0.595/0.309) | A737 | A737* |
| | - | - | - | - | - | S743 | S742 | T738 | T738 |
| | - | - | - | - | - | E745 | E744 | E740 | A740* |
| | - | - | - | - | - | P746 | A745/T745 (0.428/0.396) | A741 | T741* |

Table S7. List NCBI accession codes for the sequences used for the phylogenetic tree inference.

| ncbi accession code | species | ncbi accession code | species |
|---|---|---|---|
| XP_036256283.1 | Molothrus ater | XP_019065031.1 | Fukomys damarensis |
| XP_005423579.1 | Geospiza fortis | XP_004639300.1 | Octodon degus |
| XP_005482704.1 | Zonotrichia albicollis | XP_005377288.1 | Chinchilla lanigera |
| XP_030093883.1 | Serinus canaria | XP_005319521.1 | Ictidomys tridecemlineatus |
| XP_021401801.1 | Lonchura striata domestica | XP_003469853.1 | Cavia porcellus |
| XP_038009369.1 | Motacilla alba alba | XP_008590477.1 | Galeopterus variegatus |
| XP_041261364.1 | Onychostruthus taczanowskii | XP_045420227.1 | Lemur catta |
| XP_041874339.1 | Corvus kubaryi | XP_008061008.1 | Carlito syrichta |
| XP_039579687.1 | Passer montanus | XP_008828979.1 | Nannospalax galili |
| XP_039918735.1 | Hirundo rustica | XP_044991818.1 | Jaculus jaculus |
| NXH49971.1 | Dicaeum eximium | XP_027629197.1 | Tupaia chinensis |
| XP_032912856.1 | Catharus ustulatus | XP_004676967.1 | Condylura cristata |
| XP_005039223.1 | Ficedula albicollis | XP_048195806.1 | Perognathus longimembris pacificus |
| XP_014724522.1 | Sturnus vulgaris | XP_028746752.1 | Peromyscus leucopus |
| NXA75387.1 | Thryothorus ludovicianus | XP_021017527.1 | Mus caroli |
| XP_023774723.1 | Cyanistes caeruleus | XP_004479043.1 | Dasypus novemcinctus |
| XP_027743972.1 | Empidonax traillii | XP_037693397.1 | Choloepus didactylus |
| XP_027535188.1 | Neopelma chrysocephalum | XP_006882669.1 | Elephantulus edwardii |
| XP_047924367.1 | Anser cygnoides | XP_004390776.1 | Trichechus manatus latirostris |
| XP_027315248.1 | Anas platyrhynchos | XP_007942749.1 | Orycteropus afer afer |
| NXL86920.1 | Alectura lathami | NP_001129435.1 | Sus scrofa |
| NXC50793.1 | Penelope pileata | XP_037023606.1 | Artibeus jamaicensis |
| NXJ03225.1 | Odontophorus gujanensis | AIJ04812.1 | Taphozous melanopogon |
| XP_021240219.1 | Numida meleagris | XP_036748345.1 | Manis pentadactyla |
| XP_042728216.1 | Lagopus leucura | XP_060229914.1 | Meriones unguiculatus |
| XP_029871101.1 | Aquila chrysaetos chrysaetos | ADE75006.1 | Murina leucogaster |
| XP_026696926.1 | Athene cunicularia | XP_004320315.1 | Tursiops truncatus |
| XP_009813276.1 | Gavia stellata | EHB08960.1 | Heterocephalus glaber |
| XP_009570914.1 | Fulmarus glacialis | XP_042540604.1 | Dipodomys spectabilis |
| XP_042648655.1 | Tyto alba | XP_004602180.1 | Sorex araneus |

### Continuation of Table S7

| ncbi accession code | species | ncbi accession code | species |
|---|---|---|---|
| XP_009326612.1 | Pygoscelis adeliae | XP_037367109.1 | Talpa occidentalis |
| XP_009461166.1 | Nipponia nippon | XP_004702662.1 | Echinops telfairi |
| XP_009693986.1 | Cariama cristata | NP_001135733.1 | Oryctolagus cuniculus |
| XP_009562019.1 | Cuculus canorus | NP_945350.1 | Homo sapiens |
| XP_010086019.1 | Pterocles gutturalis | XP_045243469.1 | Macaca fascicularis |
| XP_009986007.1 | Tauraco erythrolophus | XP_004591411.1 | Ochotona princeps |
| XP_009481643.1 | Pelecanus crispus | XP_040827316.1 | Ochotona curzoniae |
| XP_009945683.1 | Leptosomus discolor | KAB1276031.1 | Camelus dromedarius |
| XP_037246269.1 | Falco rusticolus | ACI02071.1 | Rhinolophus ferrumequinum |
| XP_009881210.1 | Charadrius vociferus | XP_032944790.1 | Rhinolophus ferrumequinum |
| XP_009639677.1 | Egretta garzetta | AJF40182.1 | Rhinolophus ferrumequinum |
| XP_010297045.1 | Balearica regulorum gibbericeps | ACI02074.1 | Hipposideros armiger |
| XP_009869992.1 | Apaloderma vittatum | ACI02079.1 | Rhinolophus luctus |
| KFZ67571.1 | Podiceps cristatus | ACI02077.1 | Cynopterus sphinx |
| XP_010199933.1 | Colius striatus | XP_006925450.1 | Pteropus alecto |
| XP_010002455.1 | Chaetura pelagica | ADE75004.1 | Eonycteris spelaea |
| XP_008947217.1 | Merops nubicus | ACI02076.1 | Rousettus leschenaultii |
| XP_010129337.1 | Buceros rhinoceros silvestris | AEQ27771.1 | Leptonycteris yerbabuenae |
| XP_009941858.1 | Opisthocomus hoazin | ADO14483.1 | Kogia breviceps |
| XP_010159229.1 | Eurypyga helias | CAF5187026.1 | Ranitomeya imitator |
| XP_014819266.1 | Calidris pugnax | ADE75013.1 | Physeter catodon |
| XP_010183410.1 | Mesitornis unicolor | ADO14481.1 | Mesoplodon densirostris |
| XP_009076534.1 | Acanthisitta chloris | ADE75008.1 | Balaenoptera physalus |
| XP_009900559.1 | Dryobates pubescens | XP_036912978.1 | Sturnira hondurensis |
| KQK74720.1 | Amazona aestiva | XP_024413218.1 | Desmodus rotundus |
| KGL74759.1 | Tinamus guttatus | XP_036102144.1 | Molossus molossus |
| XP_005148456.2 | Melopsittacus undulatus | XP_016075078.1 | Miniopterus natalensis |
| XP_030336665.1 | Strigops habroptila | ADE75007.1 | Rhinopoma hardwickii |
| XP_025953428.1 | Dromaius novaehollandiae | XP_036285551.1 | Pipistrellus kuhlii |
| XP_013799723.1 | Apteryx mantelli mantelli | XP_008148033.1 | Eptesicus fuscus |
| XP_030310668.1 | Calypte anna | ADE75005.1 | Megaderma lyra |

Continuation of Table S7

| ncbi accession code | species | ncbi accession code | species |
|---|---|---|---|
| XP_021152991.1 | Columba livia | AEQ27774.1 | Pteronotus parnellii |
| XP_046763026.1 | Gallus gallus | XP_005862958.1 | Myotis brandtii |
| XP_003201941.2 | Meleagris gallopavo | AEQ27772.1 | Mormoops megalophylla |
| XP_015739933.1 | Coturnix japonica | ADE75011.1 | Megaptera novaeangliae |
| XP_025902176.1 | Nothoprocta perdicaria | XP_024593872.1 | Neophocaena asiaeorientalis asiaeorientalis |
| XP_009667752.1 | Struthio camelus australis | XP_022410127.1 | Delphinapterus leucas |
| XP_019364998.1 | Gavialis gangeticus | XP_033295445.1 | Orcinus orca |
| XP_019399134.1 | Crocodylus porosus | XP_007177376.1 | Balaenoptera acutorostrata scammoni |
| XP_006261320.1 | Alligator mississippiensis | ADO14480.1 | Hyperoodon ampullatus |
| XP_025070500.1 | Alligator sinensis | XP_007467117.1 | Lipotes vexillifer |
| XP_023964938.1 | Chrysemys picta bellii | XP_007888429.1 | Callorhinchus milii |
| XP_039373494.1 | Mauremys reevesii | XP_047193271.1 | Scophthalmus maximus |
| XP_024052722.2 | Terrapene carolina triunguis | GCC28608.1 | Chiloscyllium punctatum |
| XP_044836183.1 | Mauremys mutica | XP_039657837.1 | Perca fluviatilis |
| KAG6929882.1 | Chelydra serpentina | KAG5845660.1 | Anguilla anguilla |
| TFK06033.1 | Platysternon megacephalum | XP_042191261.1 | Callorhinchus milii |
| XP_037745122.1 | Chelonia mydas | XP_033832208.1 | Periophthalmus magnuspinnatus |
| XP_030410163.1 | Gopherus evgoodei | XP_032895760.1 | Amblyraja radiata |
| XP_048692359.1 | Caretta caretta | XP_030209716.1 | Gadus morhua |
| XP_038252898.1 | Dermochelys coriacea | XP_005995355.1 | Latimeria chalumnae |
| XP_006137601.1 | Pelodiscus sinensis | XP_023697136.1 | Paramormyrops kingsleyae |
| XP_032630412.1 | Chelonoidis abingdonii | XP_047660243.1 | Tachysurus fulvidraco |
| XP_028603177.1 | Podarcis muralis | XP_030640173.1 | Chanos chanos |
| XP_033017733.1 | Lacerta agilis | XP_018585544.2 | Scleropages formosus |
| XP_034983284.1 | Zootoca vivipara | XP_028306182.1 | Gouania willdenowi |
| XP_028603181.1 | Podarcis muralis | BAE75795.1 | Takifugu obscurus |
| XP_020645638.1 | Pogona vitticeps | XP_042578750.1 | Cyprinus carpio |
| XP_044279152.1 | Varanus komodoensis | XP_041110887.1 | Polyodon spathula |
| XP_008109741.1 | Anolis carolinensis | XP_013884866.1 | Austrofundulus limnaeus |
| XP_042323185.1 | Sceloporus undulatus | XP_023651296.1 | Paramormyrops kingsleyae |
| XP_015270544.1 | Gekko japonicus | XP_026882432.2 | Electrophorus electricus |

Continuation of Table S7

| ncbi accession code | species | ncbi accession code | species |
|---|---|---|---|
| XP_048357987.1 | Sphaerodactylus townsendi | XP_038667500.1 | Scyliorhinus canicula |
| XP_032077595.1 | Thamnophis elegans | XP_030220884.1 | Gadus morhua |
| XP_026557454.1 | Pseudonaja textilis | XP_034019476.1 | Thalassophryne amazonica |
| XP_026532999.1 | Notechis scutatus | XP_040028743.1 | Gasterosteus aculeatus aculeatus |
| XP_034289543.1 | Pantherophis guttatus | KAF6725554.1 | Oryzias melastigma |
| XP_039219698.1 | Crotalus tigris | XP_030628249.1 | Chanos chanos |
| XP_015666174.1 | Protobothrops mucrosquamatus | XP_015208044.1 | Lepisosteus oculatus |
| XP_025021645.1 | Python bivittatus | XP_020456913.1 | Monopterus albus |
| XP_029471991.1 | Rhinatrema bivittatum | XP_037099016.1 | Syngnathus acus |
| XP_033814420.1 | Geotrypetes seraphini | XP_010885587.2 | Esox lucius |
| XP_030071983.1 | Microcaecilia unicolor | XP_007577439.1 | Poecilia formosa |
| XP_040199424.1 | Rana temporaria | XP_029974339.1 | Salarias fasciatus |
| XP_040268475.1 | Bufo bufo | XP_028810143.1 | Denticeps clupeoides |
| XP_044136099.1 | Bufo gargarizans | XP_030575683.1 | Archocentrus centrarchus |
| KAG8577153.1 | Engystomops pustulosus | XP_018602967.2 | Scleropages formosus |
| KAG8439951.1 | Hymenochirus boettgeri | KAG9280146.1 | Astyanax mexicanus |
| XP_002933151.3 | Xenopus tropicalis | XP_029901453.1 | Myripristis murdjan |
| XP_018111261.1 | Xenopus laevis | KAG7455842.1 | Megalops atlanticus |
| XP_018108161.1 | Xenopus laevis | XP_032363930.1 | Etheostoma spectabile |
| XP_007504130.1 | Monodelphis domestica | KAF4079897.1 | Ameiurus melas |
| XP_038597097.1 | Tachyglossus aculeatus | XP_028279308.1 | Parambassis ranga |
| XP_044535659.1 | Gracilinanus agilis | XP_020785137.1 | Boleophthalmus pectinirostris |
| XP_027694688.1 | Vombatus ursinus | XP_043877147.1 | Solea senegalensis |
| XP_036615877.1 | Trichosurus vulpecula | XP_029017915.1 | Betta splendens |
| XP_043823103.1 | Dromiciops gliroides | XP_029912187.1 | Myripristis murdjan |
| XP_025305222.1 | Canis lupus dingo | XP_034529019.1 | Notolabrus celidotus |
| XP_035923360.1 | Halichoerus grypus | XP_046878718.1 | Hypomesus transpacificus |
| XP_045876718.1 | Meles meles | XP_035771674.1 | Neolamprologus brichardi |
| XP_039095034.1 | Hyaena hyaena | XP_034382164.1 | Cyclopterus lumpus |
| XP_046525804.1 | Equus quagga | XP_047245584.1 | Girardinichthys multiradiatus |
| XP_004439369.1 | Ceratotherium simum simum | XP_026183039.1 | Mastacembelus armatus |
| XP_035003799.1 | Hippoglossus stenolepis | XP_047431607.1 | Mugil cephalus |
